# Capturing self-renewing multipotent neural crest stem cells from human pluripotent stem cells

**DOI:** 10.64898/2026.01.18.700209

**Authors:** Yayoi Toyooka, Nami Kawaraichi, Daisuke Kamiya, Teruyoshi Yamashita, Yusaku Komoike, Kimiko Fukuda, Teppei Akaboshi, Hirokazu Matsumoto, Makoto Ikeya

## Abstract

During embryonic development, neural crest cells (NCCs) represent a multipotent population characterized by an inherently transient nature, rapidly differentiating into various lineages. This instability has presented a fundamental challenge, as it is exceedingly difficult to maintain these cells in a stable, multipotent state in vitro. In this study, we report a robust culture system dependent on Wnt and FGF signaling that enables the long-term (>6 months) expansion of human iPSC-derived neural crest stem cells (NCSCs). These NCSCs retain their self-renewal and differentiation capacity, validated at the single-cell clonal level. ATAC-seq analysis indicated that posterior NCSCs maintain a more permissive chromatin structure at neuronal gene loci. Furthermore, ChIP-seq analysis revealed that the key transcription factor SOX10 binds to the regulatory regions of genes involved in both maintenance and differentiation. This system provides a stable source of human NCSCs, offering a valuable platform for developmental biology, disease modeling, and regenerative medicine.

## Introduction

During embryogenesis, several types of transient stem cell populations emerge, contribute to tissue morphogenesis, and subsequently disappear. Neural crest cells (NCCs) are transient cell populations with multipotent stem-cell features. NCCs arise from the neural plate border between the neural plate and non-neural ectoderm as the neural tube closes, delaminate, and migrate to various tissues in the embryo and differentiate into multiple cell types, such as peripheral neurons, glia, melanocytes, smooth muscle cells, osteocytes, chondrocytes, and adipocytes.^1, 2^ Tracing experiments on avian NCCs using vital dye microinjection^3, 4^ and clonal analysis *in vitro*^5–7^ have shown that at least some NCC populations are multipotent. A previous study revealed that NCCs partially propagated and generated a multipotent progeny by serial cloning of rat NCCs, referring to such a multipotent population as “neural crest stem cells (NCSCs)”.^8^ This concept was aligned with the results of experiments utilizing modern techniques, such as a lineage-tracing experiment using R26R-Confetti mice, which showed that pre-migratory and migratory NCCs contained a multipotent population.^9^

Previous studies have attempted to promote the self-renewal of NCSCs isolated from embryos *in vitro* and have succeeded in maintaining NCSCs for several weeks or more extended periods^10–14^; however, no previous study has succeeded in achieving stable expansion of NCSCs that retain multipotency over a period of many months. Recently, several research groups have developed culture conditions under which NCCs can be induced from mouse and human pluripotent stem cells (hPSCs), such as human embryonic stem cells (hESCs) and human induced pluripotent stem cells (hiPSCs).^15–17^ Human NCCs can be derived from hESCs and hiPSCs by combining the suppression of transforming growth factor β (TGF-β) and the activation of the wingless/integrated (Wnt) signaling pathway, enabling them to differentiate into various NCC derivatives.^15–17^ However, they appear transiently in culture and immediately differentiate. Some approaches have successfully propagated hPSC-derived NCCs for several weeks or months^15–17^, but their differentiation into multiple cell lineages and stable expression of the multipotent marker gene *SOX10*, a gene encoding Sry-related HMG box protein 10, after long-term propagation were not demonstrated in these reports.

In this study, we established a culture system that enabled the expansion and long-term maintenance (over 6 months) of hiPSC-derived NCSCs by activating Wnt and fibroblast growth factor 2 (FGF2) signaling pathways. We also established posteriorized hiPSC-derived NCSCs (pNCSCs) under the same culture conditions. To validate this system, we performed comprehensive genetic and epigenetic analyses and assessed the differentiation potential of these cells into multiple neural crest derivatives. We present these data to demonstrate the versatility of our platform for facilitating both applied and basic research.

## Results

### Long-term expansion and multipotency of NCSCs

To establish a culture system for human NCSCs, we first aimed to maintain long-term SOX10 expression. Our previous attempts to expand hiPSC-derived NCCs using adherent culture with a medium containing EGF, FGF2, and SB431542 (SB) resulted in cell proliferation, but a rapid downregulation of SOX10 expression.^18^ Therefore, we hypothesized that non-adherent culture and additional Wnt signaling, which are important for the initial induction of NCC, might support their maintenance. To easily monitor SOX10 expression, the 201B7-SOX10-NL (SOX10-NL) iPSC line, in which the Venus (a GFP variant) fluorescent reporter gene is knocked into the 3′ UTR of the SOX10 locus of 201B7 iPSCs^19^, was used. Mini-screening by adding CHIR99021 (CHIR), a Wnt activator, to the EGF-, FGF2-, and SB-containing chemically defined medium (CDMi) and shifting from adherent to a 3D suspension culture system revealed that CDMi containing CHIR, EGF, FGF2, and SB (CEFS condition) could robustly maintain the SOX10-positive cell population over 6 months (Fig. 1a, b). The cells proliferated robustly under these conditions (Fig. 1c), maintaining high GFP expression for over 20 passages (140 days) (Fig. 1d). The population doubling time was approximately 48 h, demonstrating their stable and high proliferative capacity. After long-term culture (112 days, 16 passages), the cells consistently expressed NCC markers, such as *SOX10*, *PAX3*, *NGFR*, *TFAP2A, CDH6,* and *SNAI2*, at a level comparable to that of newly induced NCCs, but showed little to no expression of PSC markers, such as *POU5F1* and *NANOG* (Fig. 1e). These results were confirmed using other iPSC lines, 1231A3-SOX10-tdTomato (SOX10-tdT) iPSCs, in which the *tdTomato* fluorescent reporter was knocked into the 3′ UTR of the *SOX10* locus of 1231A1 iPSCs^20^ after 126 days of culture (18 passages) (Supplementary Fig. 1a–e), and 1231A3-GAPDH-tdTomato (GAPDH-tdT) iPSCs, in which the *tdTomato* fluorescent reporter was knocked into the 3′ UTR of the *GAPDH* locus of 1231A3 iPSCs^21^ after 371 days of culture (53 passages) (Supplementary Fig. 2a, b). Principal component analysis (PCA) and correlation matrix analysis using bulk RNA-sequencing (RNA-seq) revealed that the transcriptome of long-term cultured NCSCs was similar to that of freshly induced NCCs (induction day 10) and distinct from that of the original iPSCs and differentiated neurons (Fig. 1f, g). We also confirmed that the karyotypes were normal, the same as their parent iPSC lines (Supplementary Fig. 2c,d).

**Fig. 1.**
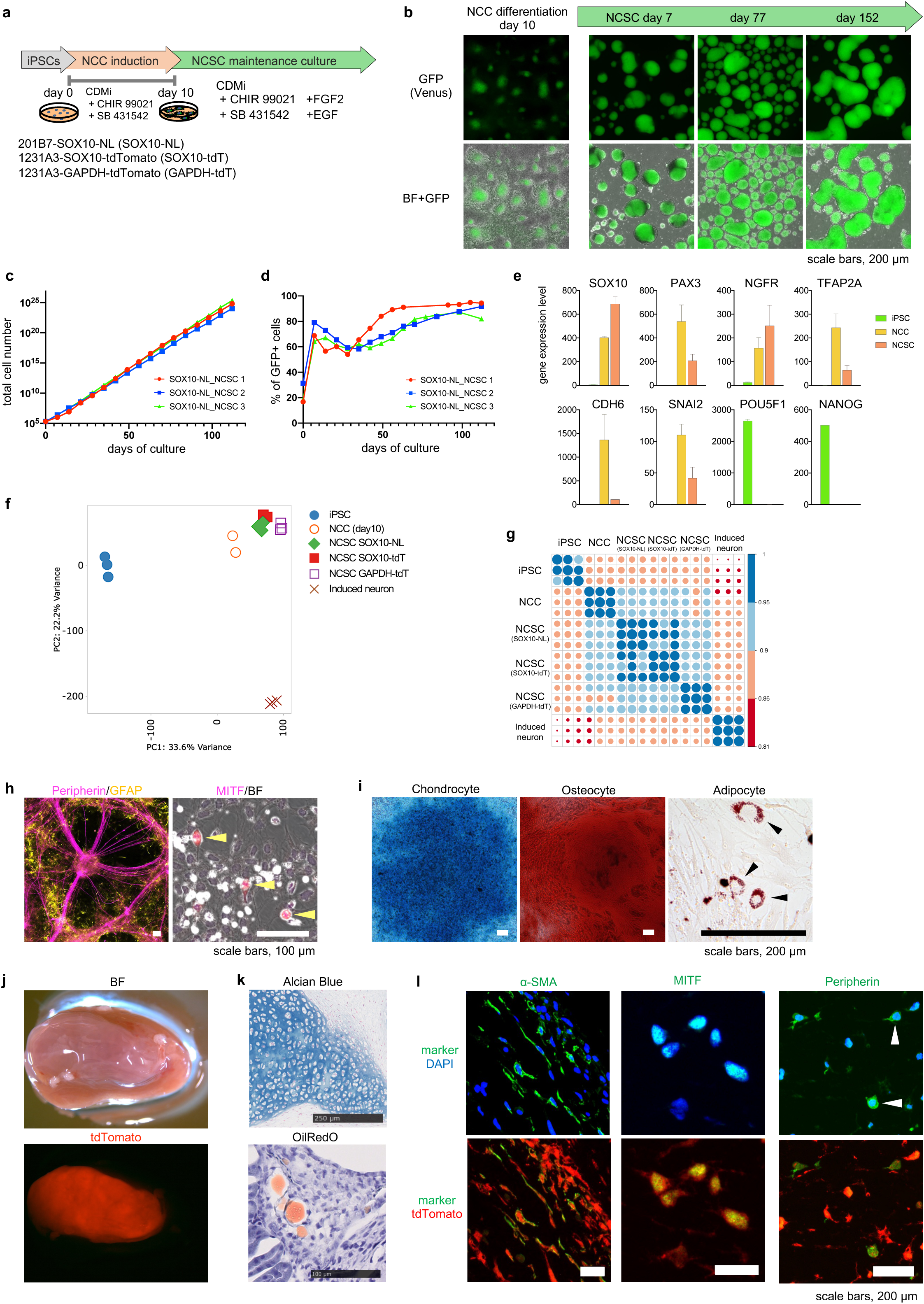
Long-term expansion and in vitro/in vivo multipotency of neural crest stem cells. **a.** Schematic procedure of the induction of neural crest cell (NCC) and maintenance as neural crest stem cells (NCSCs). **b.** Representative images of NCSCs during establishment from day 10 NCCs derived from SOX10-NL iPSCs. **c.** Growth of NCSCs (n = 3, biological replicates). **d.** Time-course changes in the SOX10+ (GFP) cell proportion in the cultures (n = 3, biological replicates). **e.** Gene expression analysis for NCC markers (*SOX10*, *PAX3*, *NGFR*, *TFAP2A*, *CDH6*, and *SNAI2*) and pluripotent cell markers (*POU5F1* and *NANOG*) in iPSCs, NCCs, and NCSCs (mean ± s.d.) (n = 3). **f, g.** PCA plot (f) and correlation matrix analysis (g) using iPSCs (SOX10-tdT, n = 3), NCCs (SOX10-tdT, n = 3), NCSCs (SOX10-NL (n = 3), SOX10-tdT (n = 3), and GAPDH-tdT (n = 3), and induced neurons (201B7, n = 3). **h, i.** Representative images of differentiated cells from long-term (84 days) maintained SOX10-NL NCSCs. **h.** Peripheral neurons (purple signals), glia (yellow signals) (left) (n = 3), and melanocytes (right, indicated by yellow arrows) (n = 3, biological replicates). **i.** Chondrocytes (left), osteocytes (middle), and adipocytes (right) (n = 3). **j.** Overall images showing bright-field (upper) and red fluorescence (tdTomato, lower) of the kidney 4 weeks after transplantation of GAPDH-tdT NCSCs (n = 3). **k.** Alcian Blue (upper) and Oil Red O (lower, indicated by black arrows) staining of sections of aggregated transplanted NCSCs (n = 3). **l.** Detection of in vivo differentiation of the transplanted NCSCs. Antibodies for α-SMA (left), MITF (middle), and Peripherin (right) were used for immunostaining. Green fluorescence indicates the signal of each lineage marker. Marker-positive cells were observed within the tdTomato+ NCSC aggregates (n = 3).

Next, we examined the differentiation potential of these long-term cultured NCSCs. When subjected to appropriate differentiation conditions, the cells (maintained for 84 days) successfully differentiated into various neural crest lineages, including peripheral neurons positive for Peripherin, glia cells positive for GFAP, melanocytes positive for MITF, and chondrocytes, osteocytes, and adipocytes stained by Alcian Blue, Alizarin Red, and Oil Red O, respectively, thereby demonstrating their multipotency *in vitro* (Fig. 1h, i). These results were also confirmed using other iPSC lines (Supplementary Figs. 1f–h, 2e). To assess their *in vivo* differentiation potential, GAPDH-tdT iPSC (ubiquitous tdTomato-labeled iPSC line)-derived NCSCs (103 days) was transplanted under the kidney capsule of immunodeficient mice. After 8 weeks, the transplant contained various cell types, including Alcian Blue-positive chondrocytes, Oil Red O-positive adipocytes, αSMA-positive smooth muscle cells, MITF-positive melanocytes, and Peripherin-positive peripheral neurons, confirming their multipotency *in vivo* (Fig. 1j–l). The migration ability was tested by transplanting aggregates of SOX10-NL NCSCs into the cranial neural crest region adjacent to the neural tube of cultured chick embryos (Supplementary Fig. 3). The transplanted cells in the spheres spread from the aggregate after 24 h of incubation, demonstrating that NCSCs can migrate. Taken together, these results suggest that our culture system can maintain bona fide NCSCs in a stable, self-renewing, and multipotent state for an unprecedentedly long period.

### Identification of key signaling factors and transcriptomic characterization of long-term cultured NCSCs

To elucidate the mechanisms underlying the long-term maintenance of NCSCs, we investigated the key factors in our culture medium. We performed exclusion tests, removing each of the four factors (CHIR, EGF, FGF, SB) individually (Fig. 2a). The removal of FGF2 resulted in a drastic reduction in the number of spheres within a week (Fig. 2b-d). The removal of CHIR resulted in a reduction in the number of SOX10-tdT-positive spheres, whereas the total number of spheres was not seriously affected. The removal of EGF or SB had minimal effects on SOX10 expression and total number of spheres. These data suggest that FGF and Wnt signaling (activated by CHIR) are essential for the maintenance of NCSCs, especially FGF for cell growth and Wnt for SOX10 expression, whereas EGF and SB are not critical.

**Fig. 2.**
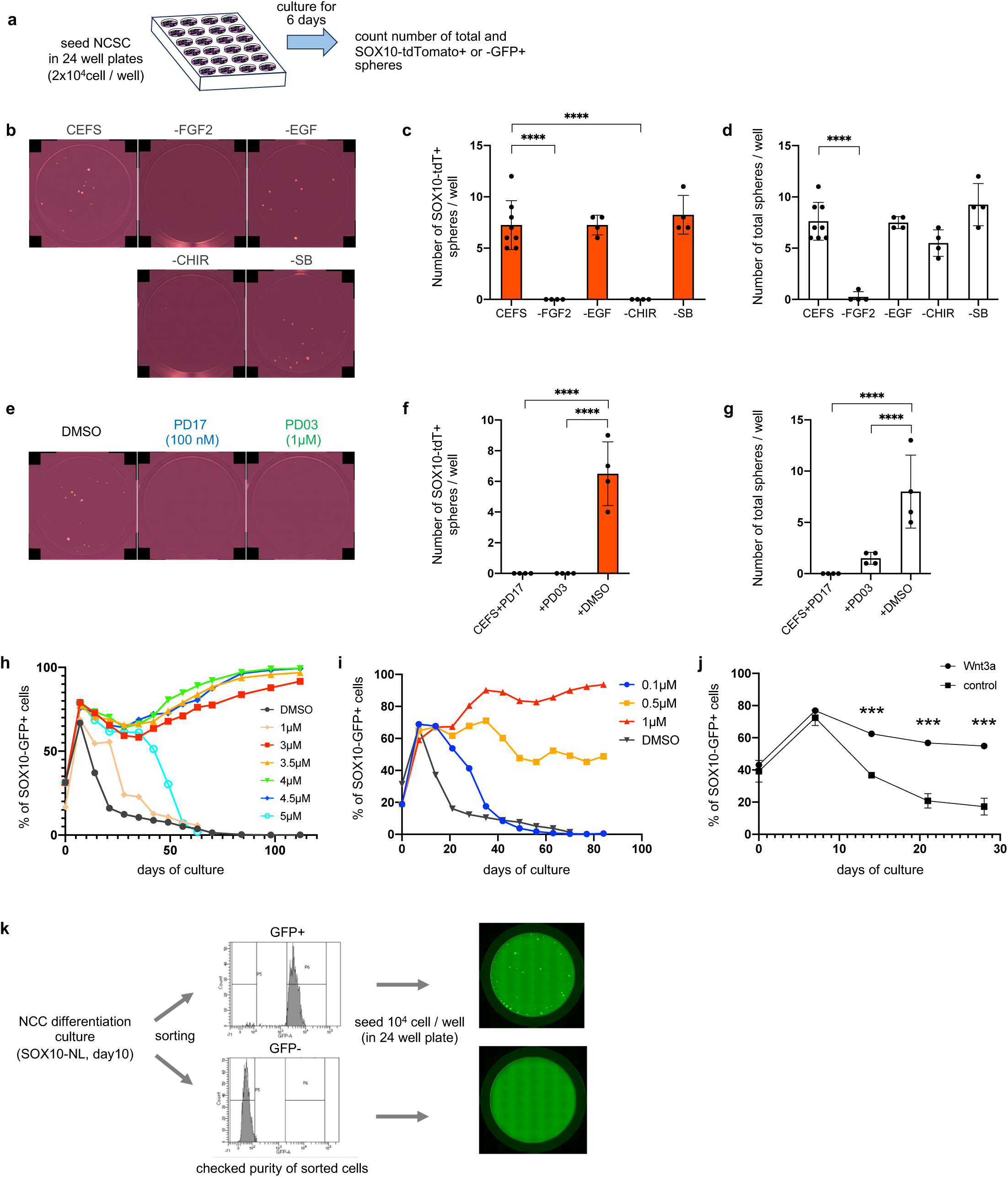
Verification of factors and signaling pathways required for the self-renewal and survival of NCSCs. **a.** Schematic overview of factor subtraction and signaling pathway inhibition experiments using SOX10-tdT NCSCs. **b.** Representative images of each well (n = 4). c, d. Graph showing the result of factor subtraction experiments for self-renewal of NCSCs. **c.** The number of SOX10+ (tdTomato+) spheres formed in the wells of a 24-well plate is shown. **d.** The total sphere number in each well is shown. Means (bars), s.d. (thin lines on bars of means), and raw data (dots) are shown. **e.** Representative images of the factor subtraction experiments (n = 4). **f, g.** Graph showing the result of inhibitor experiments for self-renewal of SOX10-tdT NCSCs. f. The number of SOX10+ (tdTomato+) spheres formed in the wells of a 24-well plate is shown. g. The total sphere number in each well is shown. Means (bars), s.d. (thin lines on bars of means), and raw data (dots) are shown. **h–j.** Maintenance experiments of SOX10+ (GFP+) NCSCs derived from SOX10-NL using the GSK inhibitors (CHIR and CP21) and human recombinant Wnt3a. Correlation between the concentration of CHIR (**h**) (n = 2 in each condition), CP21 (**i**) (n = 2 in each condition), and Wnt3a (**j**) (n = 3 in each condition) and the expansion of the SOX10+ cell population in the NCSC culture is shown. Time-course changes in the proportion of SOX10+ (GFP+) cells measured using FACS. **k.** Examination of the SOX10+ sphere formation ability of purified (sorted) GFP-positive and -negative cells in SOX10-NL NCCs on day 10 of NCC induction. P-values were calculated using a one-way ANOVA comparison of the mean values of each condition to that of the control, followed by multiple comparisons using Dunnett’s method. *P < 0.05, **P < 0.01, ***P < 0.001, and ****P < 0.0001 indicate statistical significance.

The requirement of FGF signaling was further analyzed by adding PD173074, an FGFR inhibitor, and PD0325901, an ERK inhibitor. Almost no spheres was observed in each treatment (Fig. 2e-g), suggesting that the FGF2-FGFR-ERK pathway is critical for the maintenance and proliferation of the spheres. Next, we examined the role of Wnt signaling. Since NCC induction from iPSCs requires CHIR concentrations within a narrow optimal range (0.5–1 µM), as higher (>5 µM) diverted differentiation toward alternative lineages ^17^, we treated SOX10-NL-NCCs at different concentrations of CHIR and found that 3–4.5 µM was the optimal range to maintain SOX10-NL (GFP) expression (Fig. 2h). Similar results were obtained by adding CP-21, another Wnt activator (GSK3 inhibitor), and recombinant human WNT3A (Fig. 2i, j), although a high concentration of recombinant human WNT3A (200 ng/ml) was required to maintain the SOX10+ population.

To exclude the possibility that the SOX10+ population in NCSCs was raised from the SOX10-negative (SOX10–) population contained in the culture during early passages, the SOX10+ and – fraction of SOX10-NL-NCCs (GFP+ and – at induction day 10) were sorted by GFP and cultured under the CEFS condition. The purified GFP+ population gave rise to GFP+ spheres, whereas the GFP– population formed only a small number of GFP– spheres and did not form GFP+ spheres (Fig. 2k), suggesting that only the SOX10+ population can reproduce the SOX10+ population under the CEFS condition.

### Clonal analysis confirms self-renewal and multipotency at the single-cell level

To confirm that our long-term cultured NCSCs retained the identity of the original cell population at the single-cell level, comparative single-cell RNA-sequencing (scRNA-seq) analyses were performed. The UMAP plot comparing the original NCCs at day 10 of induction after sorting and the established NCSCs after 202 days of culture (SOX10-tdT) was highly overlapped (Fig. 3a). The expression of NCC markers (*SOX10*, *PAX3*, *NGFR*, *TFAP2A*, *FOXD3*, and *CDH6*) were expressed in most major cell populations of both NCCs and NCSCs (Fig. 3b). Similar results were obtained from another NCSC line, SOX10-NL (Supplementary Fig. 4). The cell population expressing these genes was maintained at a comparative rate as the original NCCs (Fig. 3c). In contrast, the marker expression of three main cell lineages differentiated from NCCs, namely *MITF* (melanocytes), *POU4F1* (*BRN3A*, sensory neurons), and *PRRX1* (mesenchymal cells), were expressed uniformly but at low levels in most populations (Supplementary Fig. 5), suggesting that these differentiated cells were not present in NCSC cultures at high frequencies. Clustering analysis was performed using the merged data from NCCs and NCSCs, and 13 clusters were identified (Fig. 3d). Most clusters found in the original NCCs were sustained in the NCSC culture, although the ratio of subpopulations changed (Fig. 3e). Interestingly, most of the identified clusters were not characterized as differentiated cells, but suggested as different cell cycle populations (Supplementary Fig. 6a). Pseudo-time analysis revealed that the main cell cluster formed a closed loop rather than a linear trajectory (Supplementary Fig. 6b, c), which is indicative of progression through the cell cycle. Only cluster 13 showed enriched neural differentiation markers related to chemical synapses or neurotransmitter receptors, suggesting that cluster 13 contains neuronal lineage-biased or differentiating neural cells (Supplementary Fig. 6a).

**Fig. 3.**
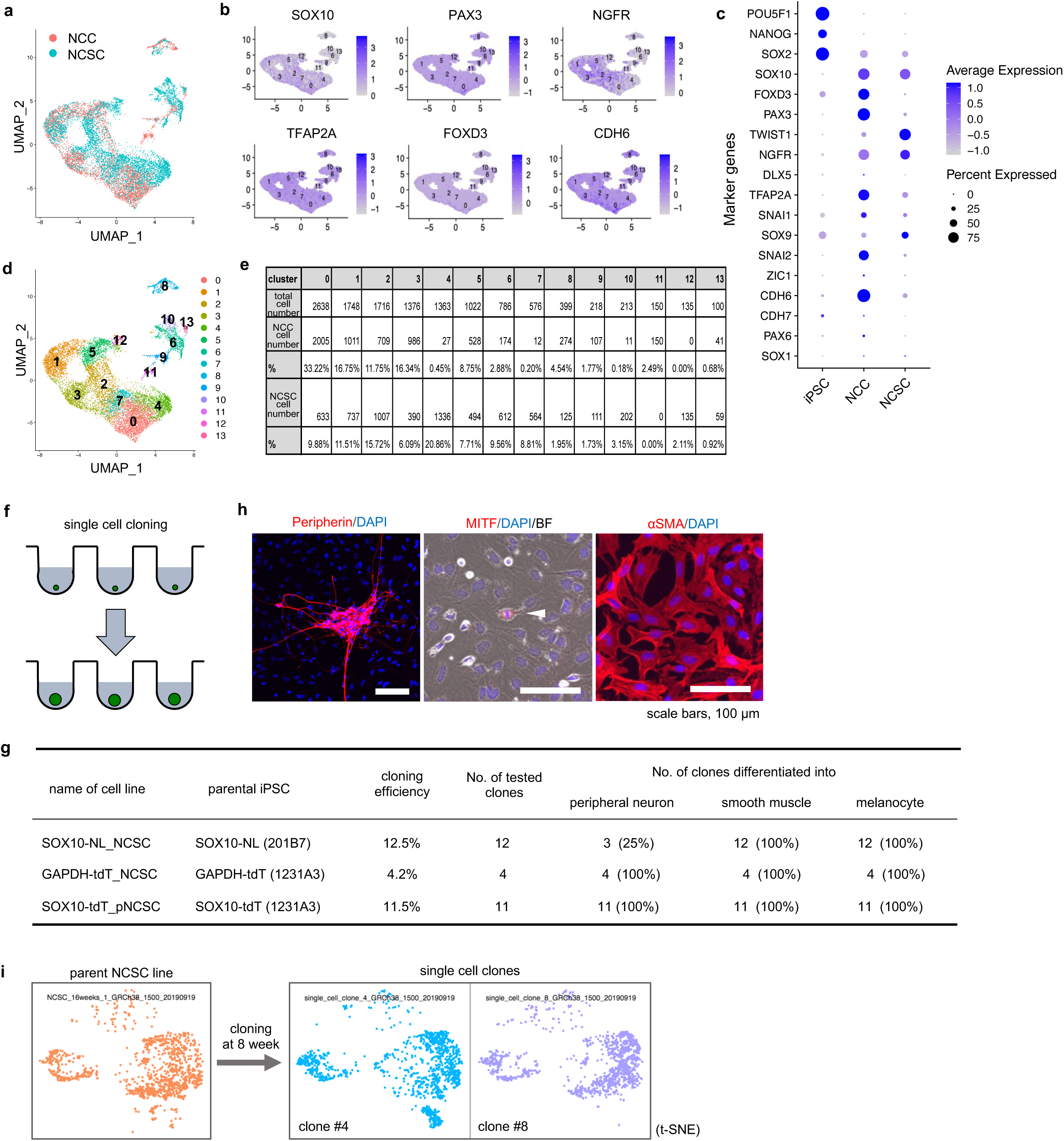
Clonal analysis confirms self-renewal and multipotency at the single-cell level. a–c. Comparison between SOX10-tdT NCSCs maintained for 202 days and original NCCs by single-cell transcriptome analysis. **a.** UMAP projection of the NCSCs and day 10 NCCs. **b.** Expression of representative neural crest marker genes (*SOX10*, *PAX3*, *NGFR*, *TFAP2A*, *FOXD3*, and *CDH6*) in the NCSCs and day 10 NCCs. **c.** Dot plot showing the expression levels of marker genes and the percentage of marker expressing cells in iPSCs, NCCs, and NCSCs. Note that expression of the pluripotent markers *POU5F1* and *NANOG*, and the markers for central nervous system *PAX6* and *SOX1* are barely detectable in original NCCs and NCSCs. **d, e.** Clusters in NCSCs (**d**) and Table (**e**) showing the cell numbers and percentages of each cluster in NCSCs and NCCs. **f.** Schematic representation of the single cell cloning and expansion. **g.** Summary of the differentiation experiments of SOX10-NL NCSC, GAPDH-tdT NCSC, and SOX10-tdT pNCSC clones. **h.** Representative images of the differentiation ability of cloned NCSCs maintained for 100 days. Cloned NCSCs were differentiated into three representative NCC lineages (neurons, melanocytes, and smooth muscle cells [SMCs]) (n = 3 in each differentiation). **i.** Comparison of the t-SNE projection of SOX10-NL NCSCs (112 days of maintenance culture) and two of its clones (56 days after cloning).

To assess the stemness of the culture at the clonal level, we next performed single-cell cloning (Fig. 3f). Single SOX10-NL+ cells were sorted into individual wells and cultured at day 100 of maintenance culture. A subset of these single cells successfully proliferated to form spheres that could be further expanded as stable cell lines (Fig. 3g). These clonally derived NCSC lines maintained high SOX10 expression and exhibited the same multipotency as the parent population, differentiating into neurons, melanocytes, and smooth muscle cells (Fig. 3g, h). Similar results were obtained from another NCSC line, GAPDH-tdT (Fig. 3g). To confirm that the clonal expansion did not alter the cellular state, we performed scRNA-seq on the established clones and compared the results with those of the parent population. The t-SNE profiles of the parent and clonal lines were nearly identical, demonstrating that the self-renewing multipotent state is a true intrinsic property of individual cells within the population and is stably inherited through cell division (Fig. 3i).

### Posterior NCSCs exhibit superior peripheral neurogenic potential

Our standard induction protocol consistently yields cranial NCCs that are negative for *HOX* gene expression.^18^ Given that NCCs exhibit distinct differentiation potentials based on their anterior-posterior positional identity^22^, it was of scientific interest to investigate whether our culture system could also support the long-term maintenance of posterior-fated NCCs (pNCCs). pNCCs were induced by the addition of retinoic acid (RA) during the induction of SOX10-tdTomato iPSCs (Supplementary Fig. 7a). Posteriorization of NCCs was confirmed by HOX gene expression (Supplementary Fig. 7b). Interestingly, pNCCs could be stably maintained and expanded in the exact same culture medium used for NCSCs, maintaining their *HOX* genes and SOX10 (tdTomato) expression (Supplementary Fig. 7b,c), multiple differentiation ability (Supplementary Fig. 7d), and similar signal requirement (Supplementary Fig. 7e–j). Single-cell cloning and induced differentiation ensured their stemness (Fig. 3g and Supplementary Fig. 7k). The reproducibility of the maintenance culture was checked with another iPSC line, 201B7 (wild-type) iPSCs (Supplementary Fig. 8).

The successful establishment of NCSCs and pNCSCs enabled us to directly compare their functional properties. Although NCSCs and pNCSCs appeared almost identical in terms of growth and sphere morphology under maintenance conditions, pNCSCs had a considerably higher neuronal differentiation potential. Comparing the neuronal differentiation of NCSCs and pNCSCs generated from the same SOX10-tdT iPSCs, a notably larger number of Peripherin-positive cells was observed in the pNCSC induction (Fig. 4a). Quantification by western blotting showed that Peripherin protein expression was approximately 30-fold higher in pNCSCs than in NCSCs after induction (Fig. 4b). These results indicate that a greater number of peripheral neurons differentiate from the pNCSC culture.

**Fig. 4.**
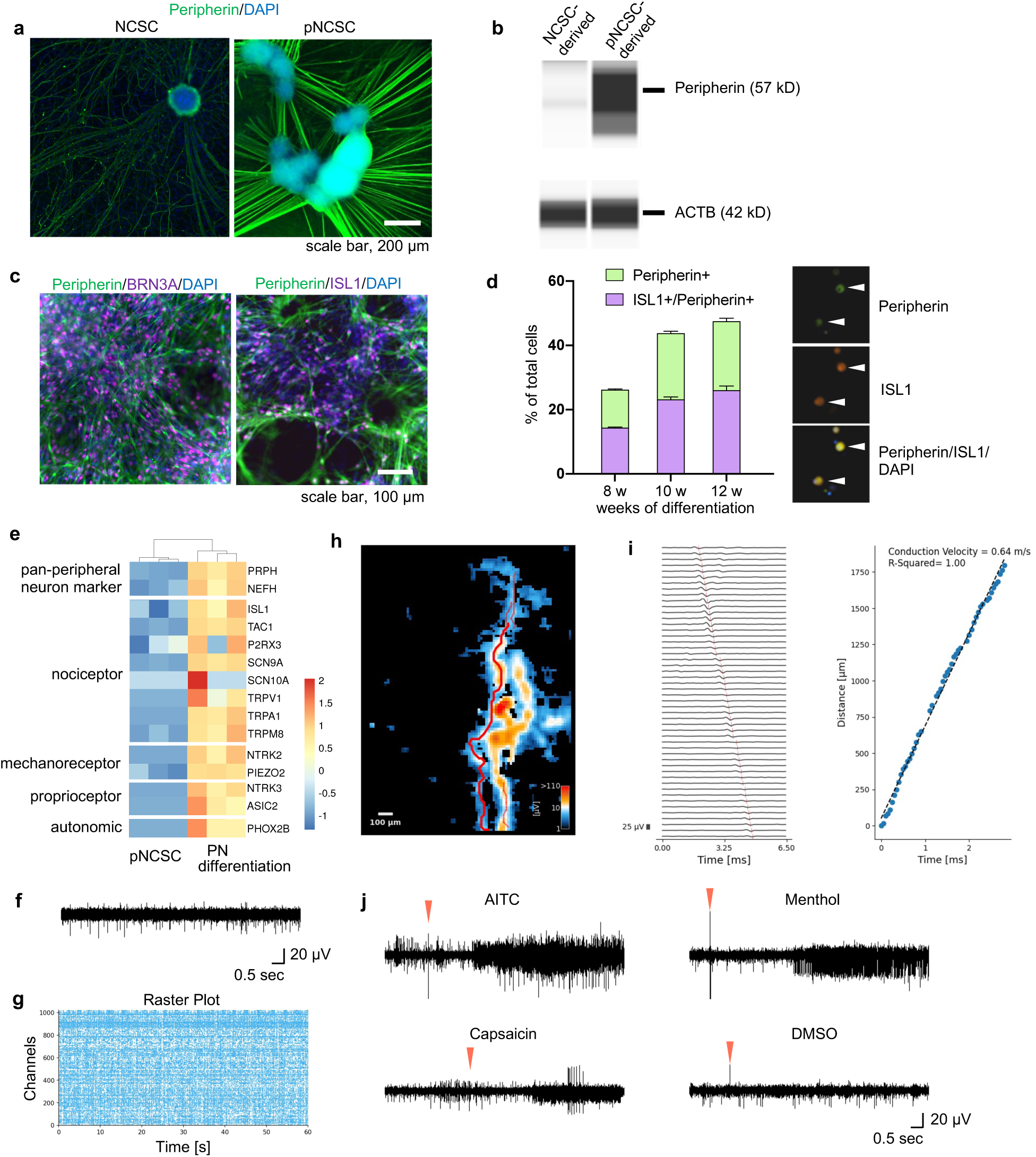
pNCSCs exhibit superior peripheral neurogenic potential. a, **b**. Comparison of the neural differentiation potential of NCSCs and pNCSCs derived from the same SOX10-tdT iPSCs. **a.** Immunocytochemical images of peripheral neurons in day 35 neural differentiation cultures from SOX10-tdT NCSCs (left) and SOX10-tdT pNCSCs (right), stained with anti-Peripherin antibody (n = 3). **b.** Comparison of Peripherin protein expression levels by western blotting after neural differentiation from NCSCs and pNCSCs (n = 3). **c.** Immunocytochemical images of peripheral neurons from 201B7 pNCSCs, stained anti-Peripherin, anti-BRN3A, and anti-ISL1 antibodies on day 14 of neural differentiation. **d.** Percentage of sensory neurons differentiated from the pNCSCs. Cultures were dissociated, adhered to glass slides, and immunostained with anti-Peripherin and anti-ISL1 antibodies (right). Positive cells were counted by image analysis (n = 4) at 8, 10, and 12 weeks (days 56, 70, and 84) in neural differentiation (left). **e.** Heatmap showing the expression of marker genes of sensory-neuron subtypes in day 84 neural differentiation of SOX10-tdT pNCSCs (n = 3, biological replicates). Expression intensities are shown as Z-scores. PN differentiation: Peripheral neuron differentiation. **f, g.** Results of MEA analysis of the pNCSC-derived peripheral neurons. **f.** Raw wave form of spontaneous firing of neurons derived from pNCSCs on day 28 of neural differentiation culture. **g.** Representative patterns of Raster plots of potentials recorded using pNCSC-derived neurons. Data on day 42 are shown (n = 3). **h, i.** Representative results of axon-tracking analysis of pNCSC-derived neurons by high density (HD)-MEA. Examples of an axon detected by axon tracking analysis (**h**); transmission of the axon potential through electrodes in the same axon shown in **h** (**i**, left); and the calculation of conduction velocity (**i**, right) are shown. Data on day 42 are shown (n = 4). **j.** Responses of pNCSC-derived sensory neurons to AITC, capsaicin, and menthol. The red arrows indicate the time at which 100 nM AITC (top left), 100 nM capsaicin (bottom left), 100 nM menthol (top right), and DMSO (control, bottom right) were administered. Data on day 70 of neural differentiation are shown (n = 4).

The culture of peripheral neurons derived from pNCSCs was confirmed to contain cells positive for the sensory neuron markers BRN3A (POU4F1) and ISL1, indicating the generation of sensory neuron-like cells (Fig. 4c). The percentage of Peripherin+/ISL1+ sensory neurons was measured by immunostaining, increasing from 14.4% at 8 weeks to 26.0% at 12 weeks (day 84) on average, demonstrating that a quarter of cells in the mature cultures were sensory neurons (Fig. 4d). To characterize this population further, RNA-seq analysis on day 84 cultures revealed that marker genes for three sensory neuron subtypes—nociceptor (e.g., *ISL1*, *TAC1*, *P2RX3*), mechanoreceptor (*NTRK2*, *PIEZO2*), and proprioceptor (*NTRK3*, *ASIC2*)—were all upregulated (Fig. 4e). Increased expression of the autonomic neuron marker *PHOX2B* was also observed.

We next performed microelectrode array (MEA) analysis to determine whether these neurons were functional (Fig. 4f–j). We observed spontaneous firing of pNCSC-derived neurons after 4 weeks of induction (Fig. 4f), although synchronized network bursts were not detected by network analysis, which is consistent with previous reports on iPSC-derived sensory neurons (Fig. 4g). An axon-tracking analysis calculated the mean conduction velocity to be 0.592 m/s, which is reasonable for unmyelinated neurons and close to reported values for rat dorsal root ganglion (DRG) sensory neurons.^23^ Finally, we examined the response of neurons to chemical stimulants. After 10 weeks of differentiation, a dramatic increase in the firing spike rate was observed immediately after the administration of 100 nM AITC, capsaicin, and menthol, thereby confirming the presence of functional sensory neurons responsive to these stimuli (Fig. 4j).

### Distinct epigenetic landscapes underlie the differential neurogenic potential of pNCSCs

Next, we investigated the epigenetic landscapes to uncover the molecular basis for the observed functional differences in neurogenic potential between NCSCs and pNCSCs. We performed an Assay for Transposase-Accessible Chromatin with sequencing (ATAC-seq) to compare their genome-wide chromatin accessibility. This analysis revealed that although key NCSC identity genes such as *SOX10* and *NGFR* showed similarly open chromatin in both cell types, the status of the loci associated with sensory neuron differentiation, *POU4F1*/*BRN3A*, *NEUROD1*, *NEUROG1*, and *NEUROG2*, was more open and accessible in pNCSCs than that in NCSCs (Fig. 5a). To identify the potential transcription factors responsible for this preconfigured landscape, we performed a de novo motif analysis using HOMER on the regions more accessible in pNCSCs was performed. This analysis revealed an enrichment for the binding motifs of key neurogenic transcription factors, including those from the POU-domain and bHLH families, at the top of the list (Fig. 5b). Consistent with this finding, bulk RNA-seq analysis confirmed that the expression of several of these key factors, including *POU4F1* (*BRN3A*), *NEUROD1*, and *NEUROG1*, was already upregulated in undifferentiated pNCSCs, whereas *NEUROG2* was not (Fig. 5c). Taken together, these results suggest that pNCSCs are maintained in an epigenetically “poised” state, which primes them for efficient differentiation toward a sensory neuron fate.

**Fig. 5.**
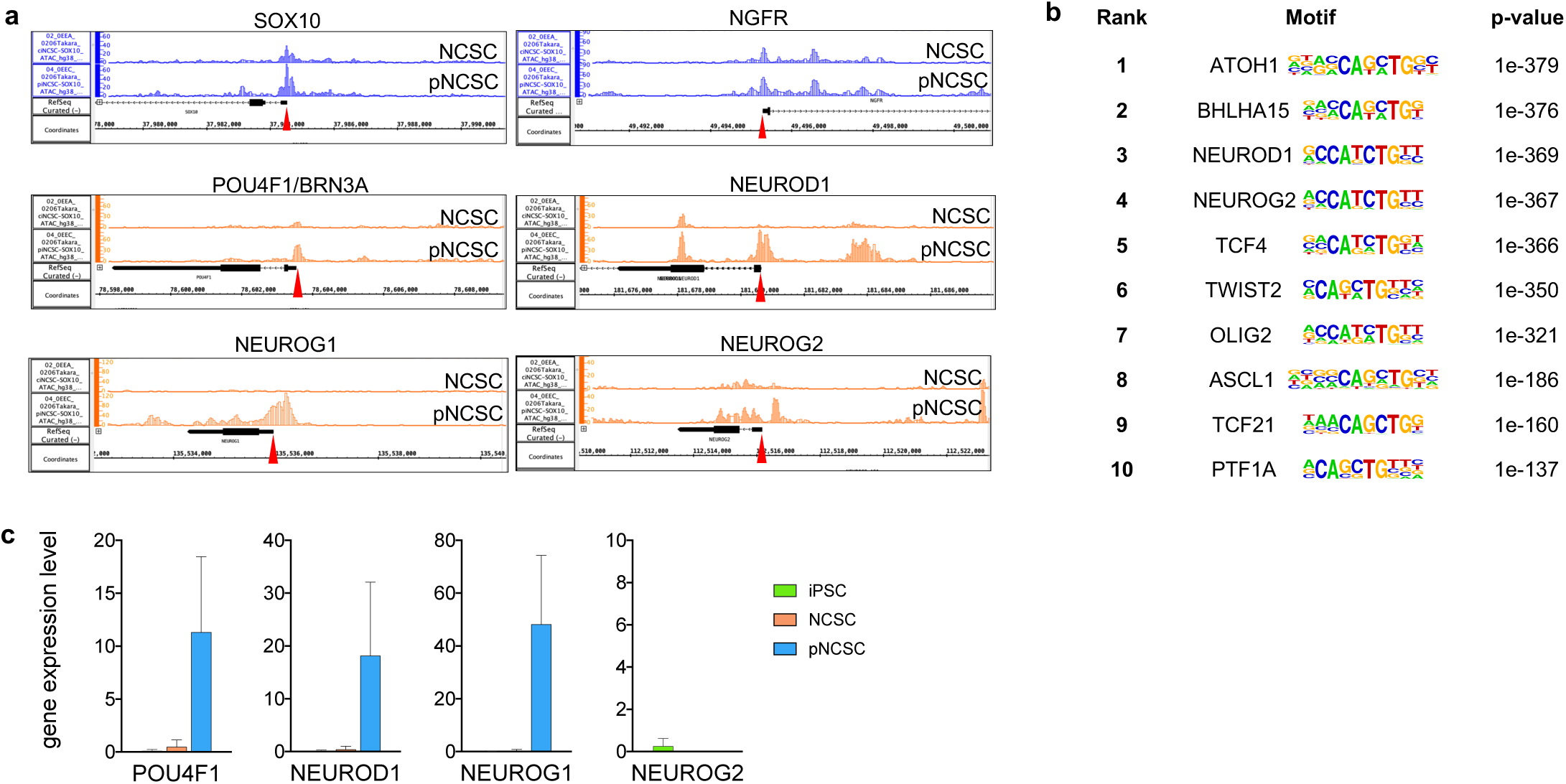
Distinct epigenetic landscapes underlie the differential neurogenic potential. **a.** Comparison of the chromatin status of loci of peripheral neuron-related genes in undifferentiated SOX10-tdT NCSCs and pNCSCs by ATAC-seq analysis. The red arrows indicate the transcription start site (TSS) of each gene. **b.** Top ten transcription factor binding motifs significantly enriched in open chromatin regions in pNCSCs compared to NCSCs identified by HOMER analysis. c. Expression of peripheral neuron-related genes (*POU4F1*, *NEUROD1* and *NEUROG2*) in undifferentiated SOX10-tdT NCSCs and pNCSCs (n = 3).

### SOX10 targets genes for both NCSC maintenance and lineage priming

Having established the epigenetic basis for the difference between NCSCs and pNCSCs, we next sought to understand the common principles that govern their self-renewal and multipotency. To this end, we investigated the role of the master transcription factor SOX10 by Chromatin immunoprecipitation (ChIP)-seq using the SOX10 antibody. We examined the gene loci in the neighboring regions of the ChIP-seq signal peak commonly detected in all (n = 4) samples and identified 1394 downstream candidate genes (Fig. 6a). Among them, we found SOX10 binding to the regulatory regions of key neural crest specifiers, including *PAX3* and *SNAI2*, as well as *SOX10* itself, suggesting a positive autoregulatory loop (Fig. 6b). Next, we performed a Gene Ontology (GO) analysis on the entire set of 1394 target genes using ShinyGO to characterize the broader biological processes regulated by SOX10 and found that terms related to neural crest development and stem cell identity, as well as to cell differentiation pathways were predominantly enriched (Fig. 6c). We also performed co-occupation analysis using the Simple Enrichment Analysis (SEA) tool in MEME Suite^24^ to identify transcription factors that bind to the neighboring regions of the SOX10 binding site. A total of 294 factors, including the neural crest-related factors FOXD3 and TFAP2B (Fig. 6d, e), were commonly identified across all samples, suggesting that these factors likely cooperate with SOX10 to regulate the expression of downstream genes. Notably, transcription factors critical for the differentiation of peripheral neurons, such as *NEUROD1*, *NEUROG2*, and *POU4F1*, were also identified (Fig. 6e). Given that both NCSCs and pNCSCs maintain their undifferentiated NCC identity in our culture system, these data suggest the bivalent nature of the NCSC state, which is simultaneously self-renewing and poised for differentiation. Additionally, the data suggest a dual role of SOX10, acting as a “gatekeeper” that simultaneously maintains the stem cell state while keeping differentiation genes primed for rapid activation upon receiving the appropriate signals (Fig. 6f).

**Fig. 6.**
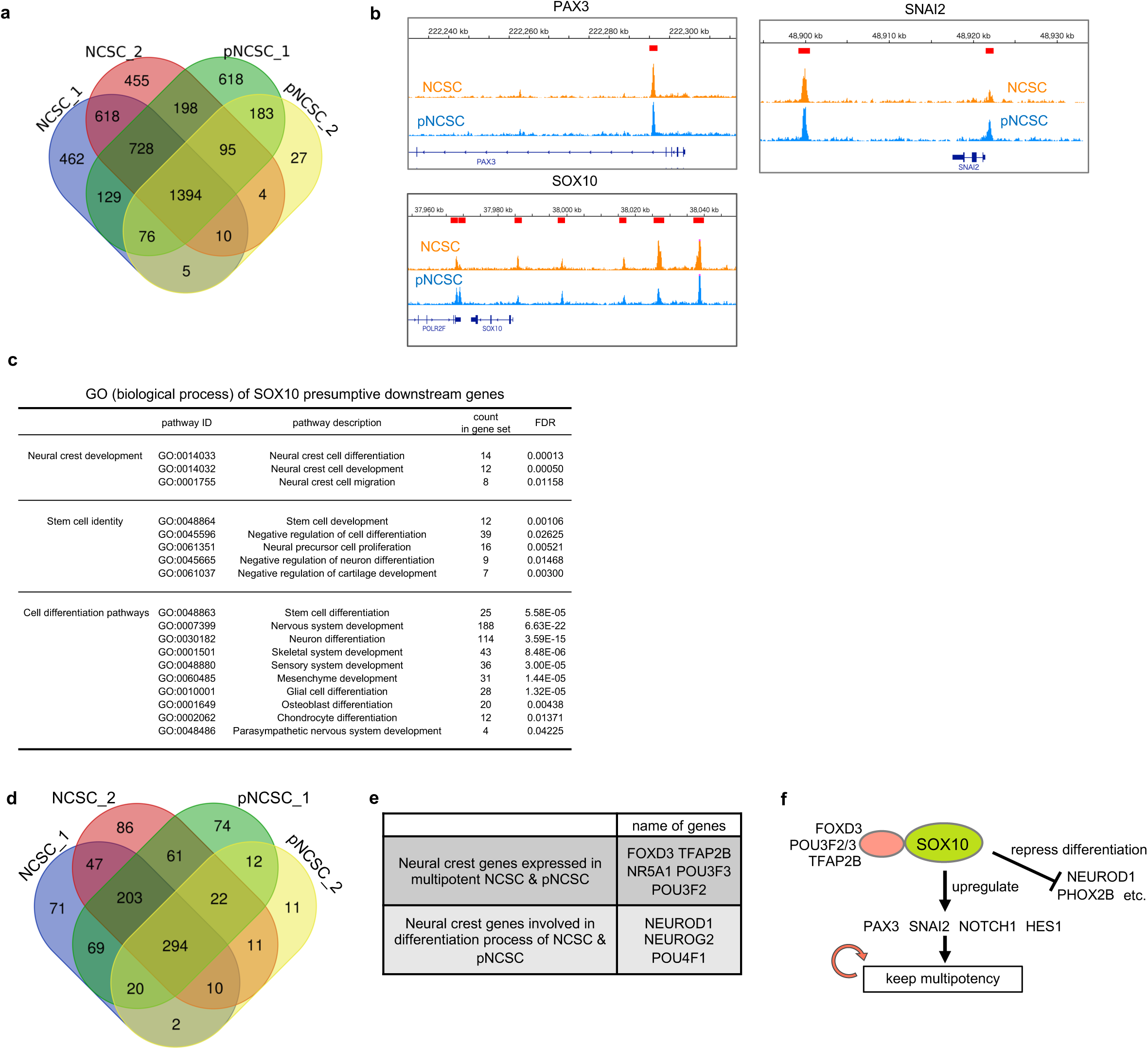
SOX10 binds to the regulatory regions of genes involved in both self-renewal and differentiation. **a–c.** Result of chromatin occupancy analysis of SOX10 ChIP-seq data using SOX10-tdT NCSCs and pNCSCs. **a.** Venn diagram of putative direct target genes of SOX10 identified by occupancy in NCSCs (n = 2) and pNCSCs (n = 2). The number of overlapping peaks are shown. **b.** Peak data of the putative direct target genes of SOX10 identified by ChIP-seq analysis using NCSCs and pNCSCs. Red lines indicate putative enhancer regions where the SOX10 peak signal is enriched. **c.** Gene Ontology (GO) processes of putative SOX10 downstream genes in NCSCs and pNCSCs. Based on a candidate gene list cut off at FDR < 0.05. **d, e.** Results of co-occupation (MEME-SEA) analysis of ChIP-seq data. **d.** Venn diagram of transcription factors identified by co-occupancy analysis using ChIP-seq data. The number of overlapping peaks are shown. **e.** Examples of neural crest-related factors cooperating with SOX10 predicted by co-occupancy analysis. f. Schematic illustration of the presumptive correlation of SOX10 and neural-crest related genes and their products in multipotent NCSCs and pNCSCs suggested from the results of ChIP-seq analysis.

## Discussion

In this study, we successfully addressed the long-standing challenge of NCC transience by establishing a culture system for the indefinite expansion of human iPSC-derived NCSCs. Our findings revealed that their lineage preference is determined by an epigenetic landscape reflecting their positional identity, and enabled us to propose a comprehensive model where these NCSCs are maintained in a poised state by the master regulator SOX10. This study provides both an unprecedented platform to interrogate human NCC development and a new framework for understanding the balance between stemness and fate specification.

The culture platform established in this study, which enables an unprecedented scale and stability in the supply of human NCSCs and pNCSCs, will make significant contributions to a wide range of fields, from basic research to clinical applications. Research into neurocristopathies, a group of disorders arising from abnormal NCC development, including neuroblastoma, Hirschsprung’s disease, and Waardenburg syndrome, has long been hampered by the difficulty of obtaining patient-derived primary cells. Our culture system overcomes this barrier by enabling an unlimited supply of patient-specific iPSC-derived NCSCs/pNCSCs, making it feasible to perform drug screening and investigate disease mechanisms at the molecular level, while also being applicable to next-generation advanced research. Although techniques to directly differentiate iPSCs into DRG organoids have recently been reported^25^, using our pNCSCs as a starting material could lead to a more stable and rapid generation of high-quality organoids, providing a more sophisticated model for studying sensory neuron development and pathology. Our system holds great potential for regenerative medicine, where a safe and large-scale supply of target cells is essential. A major advantage is that the medium is chemically defined. However, the current protocol uses bovine serum albumin (BSA), which is a hurdle that should be overcome for future clinical use. If a completely xeno-free condition (for example by substituting with recombinant albumin) can be established, this culture system will become an extremely promising foundational technology to accelerate the development of cell therapies for a variety of neural crest-related disorders.

The culture system established in this study enables us to investigate the molecular mechanisms that maintain the multipotency of NCSCs over long periods. Our ChIP-seq analysis revealed a seemingly contradictory fact: SOX10 binds to gene clusters involved in NCC development and differentiation into future lineages. The fundamental principles of stemness regulation in PSCs provide valuable insight into this dual role of SOX10. In PSC research, a strict stoichiometric balance between the transcription factors SOX2 and OCT4 functions as a “molecular seesaw” that governs the maintenance of undifferentiated states and initial fate decisions. Although these factors cooperatively activate pluripotency-associated genes, a shift in their ratio causes the cells to commit to a specific germ layer, suggesting that stem cells are poised for differentiation and exist in a dynamic equilibrium. Considering this SOX2/OCT4 paradigm, a compelling hypothesis emerges that a similar regulatory logic may exist in NCSCs. Our RNA-seq analysis confirmed that POU domain transcription factors involved in neural differentiation, such as POU3F2 (BRN2) and POU3F3 (BRN1), are expressed in long-term cultured NCSCs. In this study, we propose a model in which the fate of NCSCs is determined by a quantitative balance between SOX10 and these POU3F family factors. This hypothesis is consistent with our ChIP-seq and ATAC-seq results, which show that SOX10 is pre-bound to differentiation-related genes, possibly maintaining them in an open chromatin state. SOX10 may play a role in preparing genes for differentiation, whereas POU3F may act as the “trigger” to initiate the process. The NCSC culture system established here provides an ideal platform to test this hypothesis, such as by determining whether forced expression of POU3F2 or POU3F3 accelerates neural differentiation. Exploration of this molecular seesaw model will lead to a fundamental understanding of NCSC fate determination and will form the basis for future technologies to precisely control their differentiation.

## Methods

### Culture of human induced pluripotent stem cells

Human iPSCs (201B7, 1231A3, and their derivatives) were cultured on cell culture plates or dishes coated with iMatrix-511 (Nippi, Tokyo, Japan) in StemFit AK03N (Ajinomoto, Tokyo, Japan), as described previously.^26^

### Induction of neural crest cells (NCCs) from iPSCs

NCCs were induced as described by Fukuta et al.^17^ and Kamiya et al.^18^ For 201B7 iPSCs and their derivatives, iPSCs were seeded onto dishes coated with Matrigel (BD, Bedford, MA, USA) in mTeSR1 medium (STEMCELL Technology, Vancouver, Canada) at a density of 3.6 × 10^3^ cells/cm^2^ and cultured for 4 days. The cells were then cultured in CDMi (CDM^17^ with 7 μg/mL Insulin (FUJIFILM, Tokyo, Japan)) supplemented with 10 μM SB431542 (FUJIFILM) and 1 μM CHIR99021 (Axon Medchem, Reston, VA, USA) for 10 days to induce NCC. For 1231A3 iPSCs and their derivatives, SOX10-tdTomato and GAPDH-tdTomato, iPSCs were seeded onto iMatrix-511-coated dishes in StemFit AK03N without FGF2 at a density of 3.6 × 10^3^ cells/cm^2^ and cultured for 4 days. The cells were then cultured in StemFit Basic03 (equivalent to AK03N without FGF2, Ajinomoto) with 10 μM SB431542 and 1 μM CHIR99021 for 10 days for neural crest induction. For posteriorization, 1 μM retinoic acid (RA) was added to the induction medium from days 6 to 10 during NCC induction, with the medium changed every 2 days from days 0 to 6 and every day from days 7 to 10.

### Establishment, maintenance, and cloning of NCSCs and pNCSCs

NCCs were seeded in suspension culture dishes (SUMITOMO BAKELITE, Tokyo, Japan) at a density of 9.5 × 10^3^ cells/cm^2^ in CEFS medium (CDMi supplemented with 3 μM CHIR99021, 10 μM SB431542, 40 ng/mL EGF [FUJIFILM], and 40 ng/mL FGF2 [FUJIFILM]). Passage was performed once a week. For passaging, the cells were treated with Accutase (Innovative Cell Technologies, San Diego, CA, USA) at room temperature (20–25°C) for 3 min, neutralized by adding an equal volume of CDMi, and dissociated into single cells by pipetting. Cells were replated in suspension culture dishes at a density of 9.5 × 10^3^ cells/cm^2^. NCSC and pNCSC spheres were suspended in a GMP-grade STEM-CELL BANKER (Takara, Japan) to prepare frozen stock. NCSCs and pNCSCs were cloned by limiting dilution. Single-cell suspensions of cells were prepared and aliquoted in 96-well plates at a cell/well ratio of 1:2. The wells containing a single cell were identified via careful microscopic examination. Clones arising from single cells were expanded under the same conditions as those of the original NCSC and pNCSC lines. The cloning efficiency was calculated as the number of clones that could be expanded divided by the total number of seeded cells.

### Neural differentiation of NCSCs and pNCSCs

For peripheral neuron and glial cell differentiation, NCSC and pNCSC spheres were seeded onto fibronectin or Matrigel-coated 12-well plates and cultured in Neurobasal (Thermo Fisher Scientific, Waltham, USA) medium supplemented with B27 supplement (Thermo Fisher Scientific), N-2 supplement (Thermo Fisher Scientific), 2 mM L-glutamine (FUJIFILM, Tokyo, JAPAN), 10 ng/mL BDNF (FUJIFILM), GDNF (FUJIFILM), NT-3 (FUJIFILM), NGF (FUJIFILM), and 1% fetal calf serum for 30–45 days, with the medium changed every 3 days. Differentiated neurons were detected by immunostaining with Peripherin- and TUBB3-specific antibodies, and glial cells were detected by immunostaining with an anti-GFAP antibody.

### Differentiation of NCSCs and pNCSCs into melanocytes

For melanocyte differentiation, NCSC and pNCSC spheres were seeded onto fibronectin-coated 12-well plates and cultured in Basic03 supplemented with 1 μM CHIR99021, 25 ng/mL BMP4 (R&D Systems, Minneapolis, MN, USA), and 100 nM Endothelin-3 (TOCRIS, Bristol, UK) for 10 days, with the medium changed every 2 days. Cells were fixed on day 14, and differentiation was confirmed by immunostaining with an anti-MITF antibody.

### Differentiation of NCSCs and pNCSCs into smooth muscle cells

NCSC and pNCSC spheres were seeded onto fibronectin-coated 12-well plates and cultured in a CEFS medium. The next day, the medium was replaced with basal medium 231 (Thermo Fisher Scientific, M-231-500) supplemented with smooth muscle differentiation supplement (Thermo Fisher Scientific, S-008-5). The medium was changed every 2–3 days. The cells were fixed on day 14, and differentiation was confirmed by immunostaining with an anti-α-SMA antibody.

### Induction of mesenchymal stromal/stem cells (MSCs) from NCSCs

NCSC spheres were seeded onto fibronectin-coated plates in Basic03 medium supplemented with 10 µM SB431542, 20 ng/mL EGF, and 20 ng/mL FGF2. The next day, the medium was replaced with MSC Expansion medium (CTS StemPro MSC SFM; Thermo Fisher Scientific). The cell morphology began to change approximately 4 days after induction. Passage was performed every 4 days using Accutase. Human MSC markers (CD44, CD45, CD73, CD90, CD105, and HLA-DR) were analyzed using fluorescence-activated cell sorting (FACS) at passages 2 and 5 after MSC induction. The antibodies used for FACS analysis are listed in Supplementary Table 1.

### Differentiation of NCSC-derived MSCs

For chondrogenic differentiation, 1.5 × 10^5^ MSCs were suspended in 5 µL of chondrogenic medium (DMEM/F12, Thermo Fisher Scientific), 1% (v/v) ITS+ premix (Corning, Corning, NY, USA), 0.17 mM AA2P (Sigma-Aldrich, St. Louis, MO, USA), 0.35 mM proline (Sigma-Aldrich), 0.1 mM dexamethasone (Sigma-Aldrich), 0.15% (v/v) glucose (Sigma-Aldrich), 1 mM sodium pyruvate (Thermo Fisher Scientific), 2 mM GlutaMAX (Thermo Fisher Scientific), and 0.05 mM MTG (Sigma-Aldrich) supplemented with 40 ng/mL PDGF-BB (PeproTech, Rocky Hill, NJ, USA), 100 ng/mL TGF-β3 (R&D), 10 ng/mL BMP4, and 1% (v/v) fetal bovine serum (FBS; Thermo Fisher Scientific) and were subsequently transferred to fibronectin-coated 24-well plates. After 1 h, 1 mL of chondrogenic medium was added, and the cells were cultured for 14 days. The differentiation properties of the cells were confirmed using Alcian blue staining. Briefly, induced cells were fixed for 30 min with 4% paraformaldehyde (PFA) (FUJIFILM) and rinsed with phosphate-buffered saline (PBS). The cells were then stained with Alcian Blue solution (1% Alcian Blue, Muto Pure Chemical Co., Ltd, Tokyo, Japan) for 1 h at 25°C.

For osteogenic differentiation, 4 × 10^4^ MSCs were seeded onto 12-well plates coated with 0.1% gelatin and cultured in MSCgo Rapid Osteogenic Differentiation Medium (Biological Industries, Cromwell, CT, USA) for 30 days, with the medium changed every 3 days. Differentiation was confirmed by the formation of calcified nodules, as detected using Alizarin Red staining (Merck, Darmstadt, Germany). Briefly, culture wells were washed twice with PBS and fixed in 100% ethyl alcohol for 10 min at room temperature. Alizarin Red solution (40 mM, pH 4.2) was added to each well and incubated at room temperature for 10 min. Nonspecific staining was removed by washing several times with water.

For adipogenic differentiation, 4 × 10^4^ MSCs were seeded into a 12-well plate coated with 0.1% gelatin and cultured in hMSC Adipogenic Differentiation Medium (Lonza, Basel, Switzerland) for 32 days. The medium was changed every 3 days. Differentiation was detected using Oil Red O staining. Cells were fixed in 10% formalin for 1 h at room temperature, followed by incubation in 0.3% Oil Red O staining solution (Sigma-Aldrich) for 20 min. Nonspecific staining was eliminated by washing several times with water.

### FACS and flow cytometric analysis

FACS and flow cytometric analyses were performed using a BD FACSAria Fusion Cell Sorter (BD Biosciences, Franklin Lakes, NJ, USA) and FlowJo software (BD Biosciences, Franklin Lakes, NJ, USA) according to the manufacturer’s protocol. In all experiments, an isotype control or a control without primary antibody was used to remove nonspecific background signals. The antibodies used for FACS and flow cytometric analyses are listed in Supplementary Table 1.

### Immunocytochemistry

Prior to immunostaining with antibodies, the cells on plates were fixed with 4% PFA/PBS (FUJIFILM) at 4°C for 15 min, washed twice with PBS, and incubated with 0.3% Triton-X100 at 4°C (as the surface-active agent for penetration processing) for 30 min. Any nonspecific binding was blocked with 3% BSA/PBS at 4°C for 1 h. The nuclei were counterstained with DAPI (1:1000; Thermo Fisher Scientific). Supplementary Table 1 lists the primary and secondary antibodies used in this study. Observation and acquisition of images of the samples were performed using a BZ-X700 (Keyence, Osaka, Japan). For the experiment to count Peripherin+ and ISL1-positive cells in immunostained neural differentiation cultures, cultures were dissociated using a papain dissociation system (Worthington Biochemical Corporation, Lakewood, NJ) according to the manufacturer’s instructions. In brief, neural cultures were washed with PBS, incubated for 20 min at 37°C in a mixed solution containing papain (at a final concentration of 20 units/ml) and DNase (0.005%), and dissociated by gentle pipetting. Dissociated cells were collected, washed with wash buffer, filtered, cultured in Matrigel-coated 4-well slide chambers overnight, and fixed in 4% PFA for 1 h at 4°C. Immunostaining was performed using antibodies against neural markers, and DAPI (Thermo Fisher Scientific) was used for counterstaining. Images were acquired using a Nanozoomer S60 C13210-01 (Hamamatsu Photonics, Hamamatsu, Japan), and positive cells were detected and counted using HALO image analysis software (Indica Labs, Albuquerque NM, USA).

### Karyotype analysis

Karyotype analysis was performed at the chromocenter using quinacrine–Hoechst staining (Tottori, Japan). Chromosome counts were performed on 20 cells in metaphase, and karyotyping was performed on eight of these cells for each cell line.

### RNA-seq analysis

Total RNA was purified using the RNeasy Micro Kit (Qiagen, Valencia, CA, USA) and treated with the DNase-One kit (Qiagen) to remove genomic DNA. We reverse-transcribed 10 ng of total RNA using a SuperScript VILO cDNA Synthesis Kit (Thermo Fisher Scientific) to obtain single-stranded cDNA. We performed cDNA library synthesis for the Ion Ampliseq transcriptome using the Ion AmpliSeq Transcriptome Human Gene Expression Core Panel (Thermo Fisher Scientific) and the Ion Ampliseq Library Kit Plus (Thermo Fisher Scientific) according to the manufacturer’s protocol. Barcode-labeled cDNA libraries were analyzed using the Ion S5 XL System (Thermo Fisher Scientific) and the Ion 540 Chip Kit (Thermo Fisher Scientific).

### Single-cell RNA-seq analysis

Cells were suspended in 1 mL of PBS containing 5% FBS (FUJIFILM) and counted. The single-cell suspension was processed using the Chromium Controller (10x Genomics) with the Single Cell 3′ Reagent Kit (Chromium Next GEM Single Cell 3′ GEM, Library & Gel Bead Kit v3.1, catalog number: 1000121; Chromium Next GEM Chip G Single Cell Kit, catalog number: 1000120; 10x Genomics) following the manufacturer’s protocol. Subsequently, the library was sequenced using HiSeq2500 (Illumina, San Diego, CA, USA). Sequence data were processed using Cell Ranger software.

### Transplantation of NCSCs and pNCSCs in NOG mice

NOD.Cg-PrkdcscidIl2rgtm1Sug/ShiJic (NOG) mice were purchased from CIEA (Kanagawa, Japan). NCSC spheres were transplanted under the kidney capsules of 8–9-week-old male NOG mice under general anesthesia with 1.5%–2% isoflurane. The transplanted kidneys were excised at 4 or 8 weeks after the transplantation of NCSCs and were fixed overnight at 4°C with 4% PFA/PBS (FUJIFILM). All animal experiments were approved by the Institutional Animal Research Committee.

### Transplantation of NCSCs in chick embryos

Fertilized chicken eggs were purchased from local farmers and incubated at 38℃ to obtain embryos at stage 9–10 on the Hamburger–Hamilton (HH) stage.^27^ Embryos were explanted in Pannett–Compton saline^28^ using a modified version of the new culture method.^29, 30^ One sphere (aggregate) of NCSCs derived from SOX10-NL iPSCs was transplanted into the cranial neural crest region adjacent to the neural tube. The embryos were incubated for 24 h, and the migration of NCSCs was observed using a fluorescent stereomicroscope (Olympus).

### Immunohistochemistry

The fixed mouse kidneys were washed in PBS, passed through a graded series of ethanol solutions, and embedded in paraffin. Paraffin-embedded sections (5 μm) were deparaffinized, and antigen retrieval was performed in an autoclave. After three washes in TBS, the sections were incubated with the diluted primary antibody (in TBS) for 1 h at 4°C. After three washes in TBS, the sections were incubated with Alexa-conjugated secondary antibody (1:200, Thermo Fisher Scientific) for 1 h at 4°C and washed three times in TBS. The sections were mounted on a SlowFade Gold Antifade Mountant with DAPI (Thermo Fisher Scientific). Supplementary Table 1 lists the primary and secondary antibodies used in this study. Images were observed and acquired using a VS120-L100 (Olympus, Tokyo, Japan), BX51 with DP71 (Olympus), and Nanozoomer S60 C13210-01.

### ATAC-seq analysis

Cells of undifferentiated NCSCs and pNCSCs (1 × 10^5^) were used for ATAC-seq analysis. Sample processing and sequencing for ATAC-seq analysis were performed at Active Motif (Carlsbad, CA, USA). Peak data analysis and visualization were performed using IntegratedGenomeBrowser.

### Imaging and counting of SOX10+ spheres

Images of each well from 24-well plates in which SOX10-tdTomato or SOX10-NL NCSCs were seeded and incubated for 6–7 days were acquired using Cell3iMager (SCREEN Holdings, Kyoto, Japan). Bright-field and fluorescence images were analyzed using Cell3 Imager software. The number of NCSC spheres in each well was counted using bright-field images, and the fluorescence areas and intensities of each sphere were calculated using the fluorescence images.

### Microelectrode array (MEA) analysis of peripheral neurons from pNCSCs

MEA was performed according to the protocol provided by MaxWell Biosystems. The electrode areas on a MaxTwo HD-MEA 6-well plate (MaxWell Biosystems) were coated with Poly (ethyleneimine) (PEI, Sigma-Aldrich) and Geltrex (Thermo Fisher Scientific). pNCSCs were seeded at a density of 5 × 10^5^ or 1 × 10^6^ cells per well and cultured in neural induction medium. The electrical activity of pNCSC-derived neurons was measured, and axon tracking was performed using the MaxTwo microelectrode array system (MaxWell Biosystems). Drug administration experiments were performed on peripheral neurons after 10 weeks of neuronal differentiation culture using Maestro Pro (AXION BIOSYSTEMS). pNCSCs were seeded in a CytoView MEA 24 plate coated with Geltrex (Thermo Fisher Scientific) at a concentration of 5 × 10^5^ or 1 × 10^6^ cells per well and cultured in neural induction medium. The drugs were carefully added at 6 µL/well to wells containing 600 µL of medium. Recordings were taken using Axis Navigator software (AXION BIOSYSTEMS).

### Statistical analysis and processing of RNA-seq data

Statistical analyses were performed using GraphPad Prism 8 software (GraphPad Software Inc., San Diego, CA, USA). P-values were calculated using one-way analysis of variance (ANOVA), comparing the mean values of each condition with that of the control, followed by multiple comparisons with Dunnett’s method. *P < 0.05, **P < 0.01, ***P < 0.001, and ****P < 0.0001 were considered to indicate statistical significance. RNA-Seq data were analyzed and plotted using iDEP (http://bioinformatics.sdstate.edu/idep/), ClustVis (https://biit.cs.ut.ee/clustvis/), and PCA 3D Visualiser (https://prismtc.co.uk/resources/free-tools/pca-3d-visualiser). Spearman correlation coefficients were calculated using the R function ‘cor’ and visualized using the R library ‘corrplot’.

scRNA-seq data were analyzed using the R package Seurat v4.4.0. Low-quality cell data were excluded. All samples were integrated using the “anchor” method, and dimensional reduction was performed using principal component analysis (PCA). Then, a uniform manifold approximation and projection (UMAP) plot was generated using PC1–PC20. These single-cell profiles were clustered by shared nearest neighbor (SNN) modularity optimization (resolution = 0.6) and projected onto a UMAP plot. Highly expressed genes characterizing each cluster (p_val < 0.01, avg_log2FC > 0.25, pct.1 > 0.25, or pct.2 > 0.25) were identified by Seurat FindAllMarkers, and enrichment analysis (reactome pathway analysis) was performed using the R package clusterProfiler v4.8.1 based on the Reactome Pathway database (R package ReactomePA v1.44.0) and Gene Ontology (GO) database (R package org. Hs.eg.db v3.17.0). Pseud-time analysis was performed using the R package monocle3 v1.2.9.

### ChIP-seq analysis

Chromatin immunoprecipitation (ChIP) was performed using standard methods with anti-SOX10 antibodies (ab272074, Abcam). Recombinant rabbit IgG (ab210849, Abcam) was used as a control for the measurement of non-specific background. NCSCs and pNCSCs derived from SOX10-tdT iPSCs (1.0 × 10^7^ cells per sample) were collected in a 15 mL tube. The cells were centrifuged at 2,000 rpm for 5 min at 4°C and suspended in 10 mL of culture medium. Subsequently, 270 µL of 37% formaldehyde (1% final concentration) was added and rotated gently for 10 min at room temperature. 1 mL of 1.25 M glycine solution (125 mM final concentration) was added and rotated gently for 5 min at room temperature, before centrifuging at 4°C and 2000 rpm for 5 min, and removing the supernatant. The cells were suspended in 10 mL PBS (–/–) +complete EDTA-free (11873580001, Thermo Fisher Scientific), centrifuged at 4°C and 2000 rpm for 5 min, and the supernatant was removed. Fixed cells were lysed in 5 mL LB1 buffer (50 mM HEPES-KOH, pH 7.5 containing 140 mM NaCl, 1 mM EDTA, pH 8.0, 0.5% NP-40, 0.25% Triton X-100, and + complete EDTA-free), washed in 5 mL LB2 buffer (10 mM Tris-HCl, pH 8 containing 200 mM NaCl, 1 mM EDTA, pH 8.0, 0.5 mM EGTA, and + complete EDTA-free) to remove detergents, and resuspended in 3 mL LB3 buffer (LB2 buffer containing 0.1% Na-deoxycholate and 0.5% N-lauroylsarcosine). Chromatin was sonicated for 12 cycles (on 30 s, off 1 min 30 s) at a power of 75 using a Q700 Sonicator (QSonica, Newtown, CT, USA). After adding 3 mL of LB3, 1/10th of the total volume of 10% TritonX-100 was added, and 1 mL (3.3 × 10^6^ cells) was dispensed into 2 mL tubes. Cells were centrifuged at 14000 rpm for 10 min at 4°C, and the supernatant was collected in a new 15 mL tube. Magnetic bead-conjugated antibodies were prepared according to the manufacturer’s instructions using Dynabeads M-280 Sheep Anti-Rabbit IgG (11203D, Thermo Fisher Scientific), with 5 μg antibody used per reaction. The sonicated chromatin was incubated with 100 µL of normal rabbit IgG (magnetic beads-conjugated) at 4°C for 30 min. The supernatant was collected in a new 2 mL tube. Subsequently, 100 µL of anti-SOX10 Ab (magnetic beads-conjugated) or normal rabbit IgG (magnetic beads-conjugated) was added per tube and rotated at 4°C overnight. Beads were washed with 1 mL of each buffer in the following order: ChIP dilution buffer (one time, 16.7 mM Tris-HCl pH 8.0 containing 167 mM NaCl, 1 mM EDTA pH 8.0, 1.1% TritonX-100, and 0.01% SDS), wash buffer 1 (two times, 20 mM Tris-HCl pH 8.0 containing 150 mM NaCl, 2 mM EDTA pH 8.0, 1% TritonX-100, and 0.1% SDS), wash buffer 2 (two times, 20 mM Tris-HCl pH 8.0 containing 500 mM NaCl, 2 mM EDTA pH 8.0, 1% TritonX-100, and 0.1% SDS), wash buffer 3 (two times, 10 mM Tris-HCl pH 8.0 containing 250 mM NaCl, 1 mM EDTA pH 8.0, 1% NP-40, and 1% Na-deoxycholate), and TE (one time). Moreover, 1 mL of TE was added and suspended. Subsequently, a portion was divided for IP confirmation, and the remainder was transferred to a new 2-mL tube. After removing the supernatant, 100 µL of elution buffer was added, followed by 2 µL of 20 mg/mL proteinase K (26160, Thermo Fisher Scientific). Subsequently, the mixture was rotated at 42°C for 2 h, followed by overnight rotation at 65°C. The supernatant was collected after incubation at room temperature for 5 min. DNA was purified from the collected supernatant using the MinElute PCR Purification Kit (QIAGEN, 28004) according to the protocol provided with the kit.

### ChIP-seq data analysis

An Illumina Novaseq (control software: Novaseq Control Software 1.7.0) was used to sequence paired-end 50 bp on SOX10 ChIP samples. The quality control of the sequenced reads was performed using FastQC (http://www.bioinformatics.babraham.ac.uk/projects/fastqc/), and the adaptor sequences were trimmed using Cutadapt (ref) and BWA software v.0.7.5a.^31^ Alignment to a reference genome (hg38) obtained from the UCSC genome browser^32^, filtering reads in blacklisted regions in the ENCODE project^33^, and removal of PCR duplicates and multi-mapped reads were performed using SAMtools.^34^ Alignment coordinates were converted to the BED format using BEDTools v.2.17^35^ and aligned to the reference genome (hg38) from MACS2^36^, which was then used to detect the Peak (FDR < 0.01). From the Peak detection result bed file, genes with TSS within 100000 bp of the Peak were identified using ChIPpeakAnno.^37^ Co-occupation analysis was performed using MEME-SEA 38. IGV (https://igv.org/app/) was used to visualize the peaks. Venn diagrams were calculated and drawn using a webtool of Bioinformatics & Evolutionary Genomics (http://bioinformatics.psb.ugent.be/webtools/Venn/). GO analysis of putative SOX10 downstream genes was performed using ShinyGO v.0.85 (https://bioinformatics.sdstate.edu/go/). Notable GO pathways filtered with FDR < 0.05 are listed in Fig. 6c.

## Acknowledgements

We would like to acknowledge Drs. Austin Smith and Noriaki Sasai for his critical reading of the manuscript. Original 201B7 and 1231A3 iPSC lines were kindly provided from Dr. Yamanaka Laboratory, Kyoto University. We thank Dr. Sato for providing SOX10-NL iPSC lines. We are also grateful to Mr. Onishi of SCREEN Holdings Co., Ltd. for supporting the image data analysis using the Cell3iMager. We thank Dr. Takayuki Kamei for his advice on the HD-MEA analysis using MaxTwo. We thank Dr. Ikuro Suzuki and Dr. Nami Nagafuku for their advice on MEA analysis using Maestro Pro. We thank Axcelead Drug Discovery Partners, Inc. for performing the immunostaining experiments. We are also grateful to Dr. Yuji Mochiduki for performing the ChIP-seq experiment. We thank the staff of Orizuru Therapeutics Inc. for their assistance with cell culture and sample preparation. We thank Takara Bio USA, Inc. for providing the tdTomato vector.

## Declaration of generative AI and AI-assisted technologies

During the preparation of this work, the authors used Gemini to check grammar and refine sentences. After using this tool, the authors reviewed and edited the content as needed and take full responsibility for the final content of the publication.

## Author contributions

Y.T., H.M., and M.I. conceived and designed the experiments. Y.T., N.K., D.K., and T.A. performed cell culture experiments. T.Y. designed probes and supervised the generation of knock-in iPSC lines. Y.K. and K.F. performed cell transplantation experiments. Y.T. wrote the original manuscript and Y.T. and M.I reviewed and edited the manuscript.

---

**Supplementary Fig. 1.**
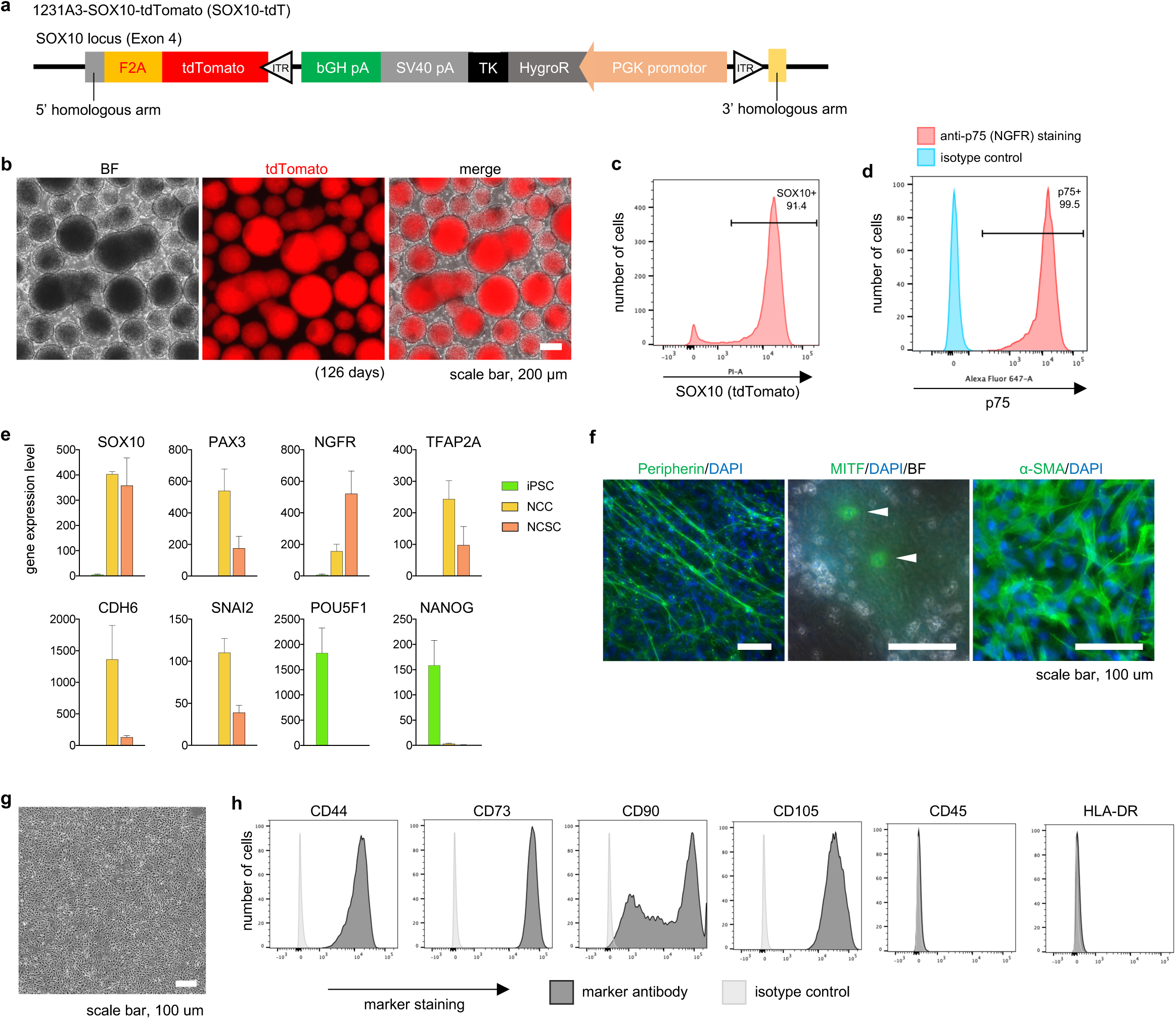
Reproducibility using 1231A3-SOX10-tdTomato iPSCs. **a.** Schematic representation of the SOX10 locus of SOX10-tdTomato (SOX10-tdT) reporter iPSC line. **b.** Morphology and tdTomato fluorescence of NCSCs established from SOX10-tdT iPSCs cultured for 126 days. Representative images of NCSC spheres. **c.** FACS analysis of SOX10-tdT NCSCs. The SOX10+ (tdTomato+) population became dominant (approximately 90%) in established NCSCs (n = 3, biological replicates). **d.** FACS analysis of p75 (NGFR) expression using anti-p75 antibody in SOX10-tdT NCSCs after long-term (112 days) expansion. **e.** Gene expression analysis of NCC markers (*SOX10*, *PAX3*, *NGFR*, *TFAP2A*, *CDH6*, and *SNAI2*) and pluripotent cell markers (*POU5F1* and *NANOG*) in iPSCs, NCCs, and NCSCs (mean ± s.d.) (n = 3). **f–h.** In vitro differentiation ability of SOX10-tdT NCSCs. **f.** Differentiation of NCSCs into peripheral neurons (left), melanocytes (middle), and smooth muscle cells (right) (n = 3). **g.** Bright field image of MSCs derived from NCSCs (n = 3). **h.** Expression of cell surface markers for MSCs (positive for CD44, CD73, CD90 and CD105, and negative for CD45 and HLA-DR) in (**g**) (n = 3).

**Supplementary Fig. 2.**
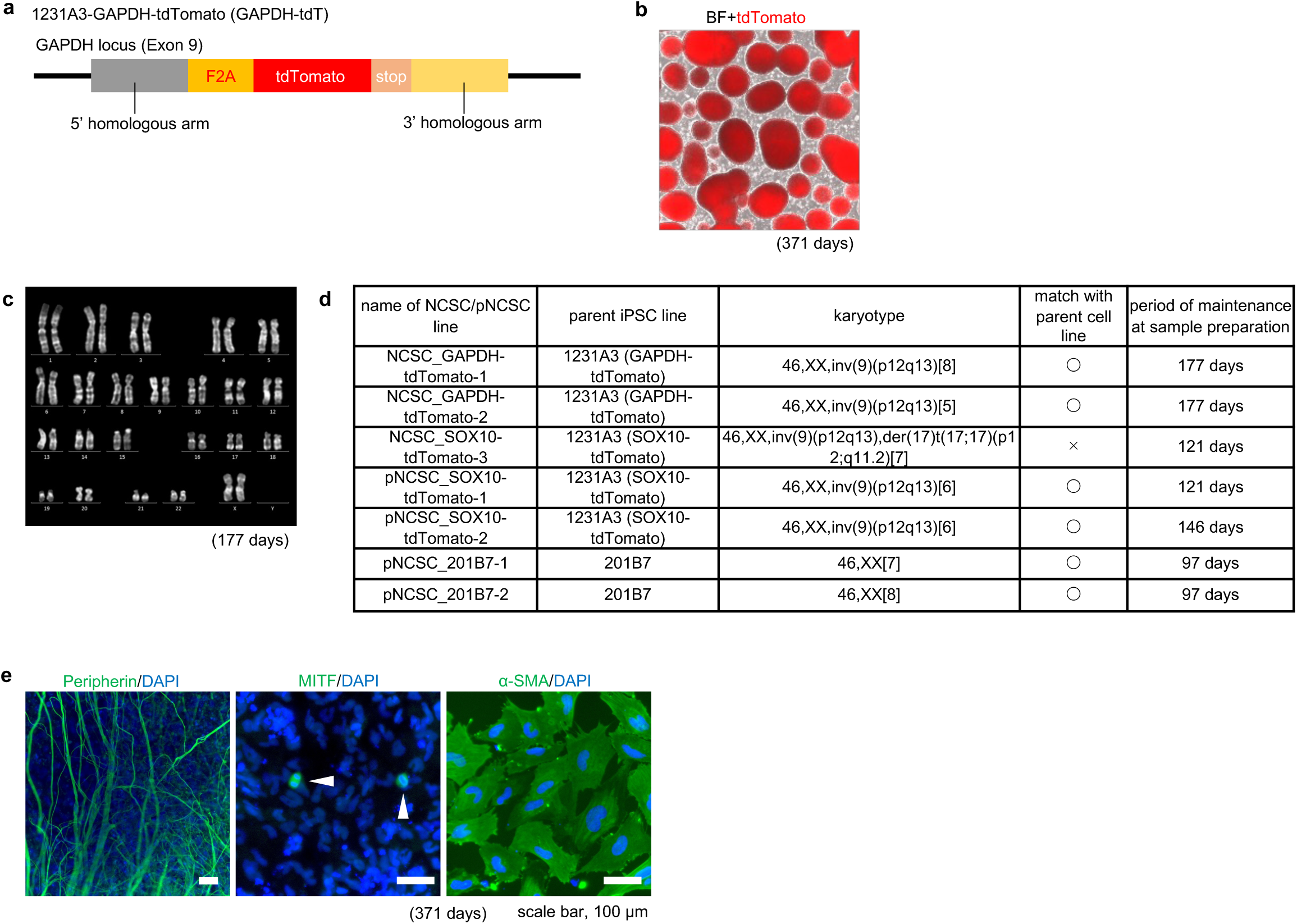
Reproducibility using 1231A3-GAPDH-tdTomato iPSCs. **a.** Schematic representation of the GAPDH locus of the GAPDH-tdTomato (GAPDH-tdT) reporter iPSC line. **b.** Morphology and tdTomato fluorescence of NCSCs established from GAPDH-tdT iPSCs cultured for 371 days. Representative images of NCSC spheres. **c, d.** Karyotype analysis of NCSCs. **c.** The result of an NCSC line derived from 1231A3 GAPDH-tdT iPSCs and maintained for 177 days is shown, proving that it has the same karyotype as the 1231A3 WT iPSCs, 46, XX, inv(9)(p12q13). Chromosome counts were performed on 20 cells in metaphase, and karyotyping was performed on eight of these cells for each cell line. **d.** Summary of the karyotype analysis for all NCSCs and posteriorized NCSCs (pNCSCs). **e.** In vitro differentiation ability of GAPDH-tdT NCSCs. Differentiation of peripheral neurons (left), melanocytes (middle), and smooth muscle cells (right) are shown (n = 3).

**Supplementary Fig. 3.**
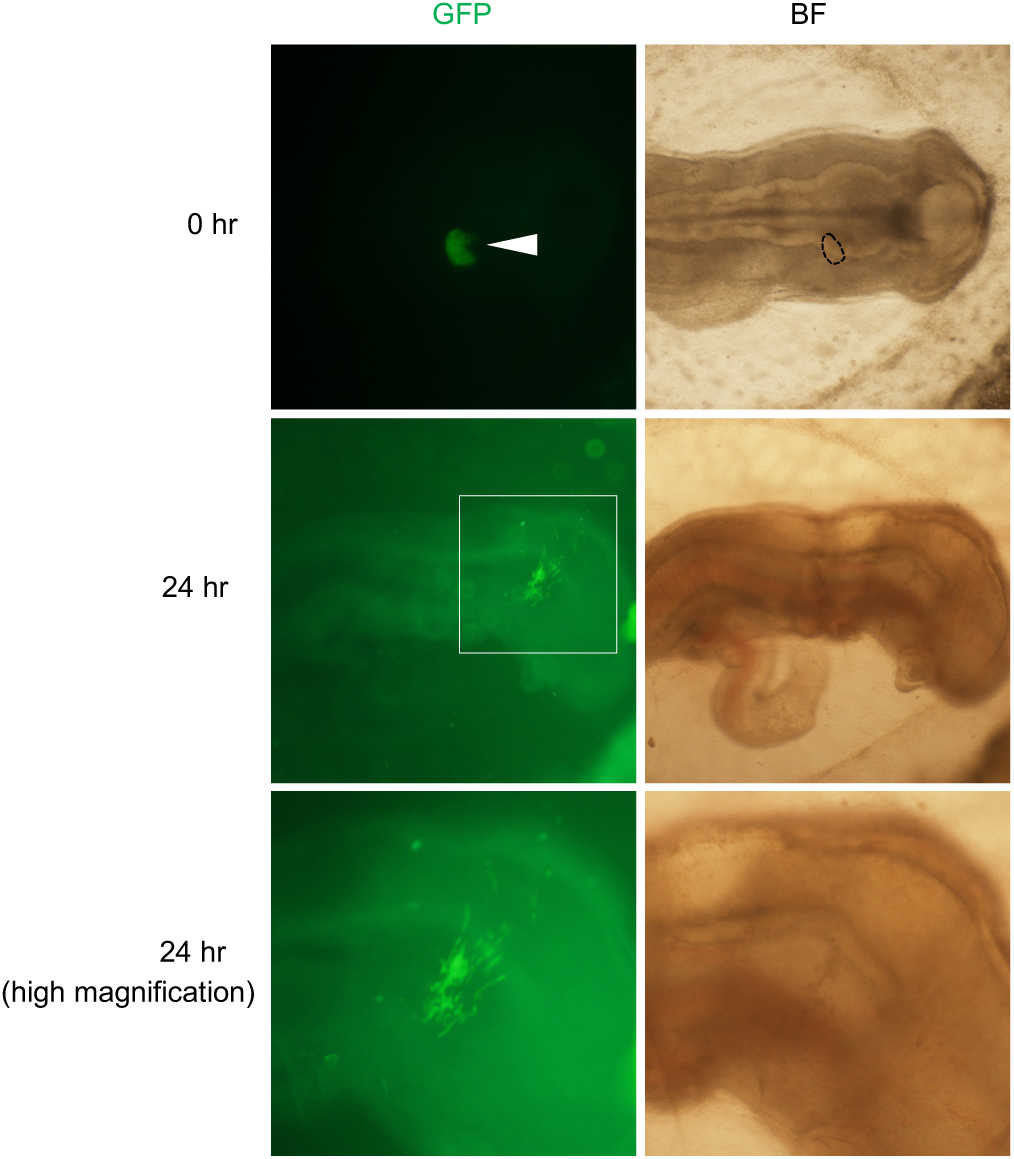
Migration assay of NCSCs in chick embryos. A sphere of SOX10-NL NCSCs expanded for 130 days was transplanted into the developing chicken embryos. Upper, middle, and lower panels show NCSCs immediately after being transplanted into the cranial neural crest region, a transplanted region of the same embryo after culture for 24 h, and high magnification of the transplanted region at 24 h. Representative results of the three independent experiments are presented.

**Supplementary Fig. 4.**
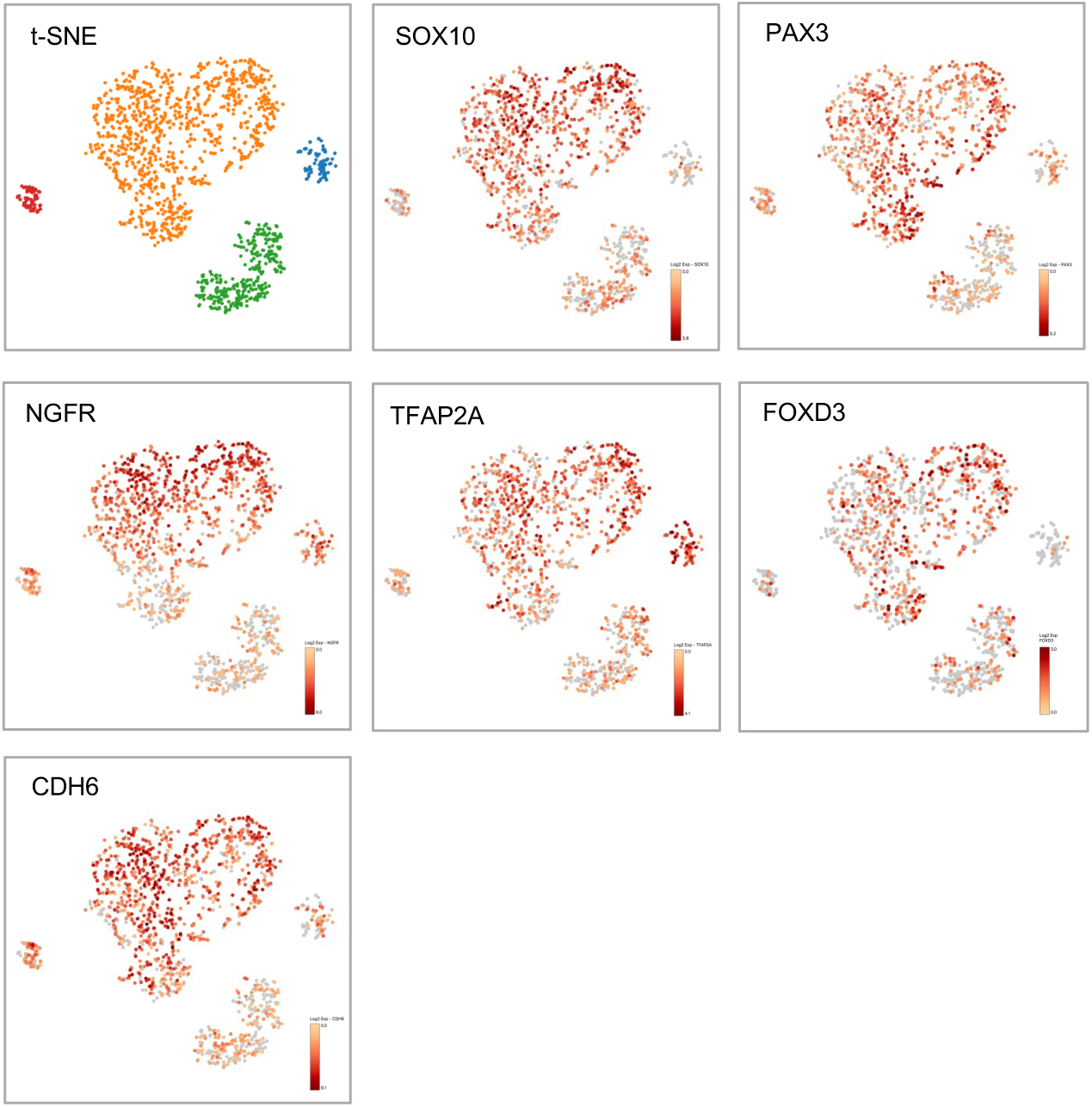
Results of single-cell transcriptome analysis of SOX10-NL NCSCs. t-SNE projection of NCSCs maintained for 112 days shows uniform expression of representative NCC markers (*SOX10*, *PAX3*, *NGFR*, *TFAP2A*, *FOXD3*, and *CDH6*).

**Supplementary Fig. 5.**
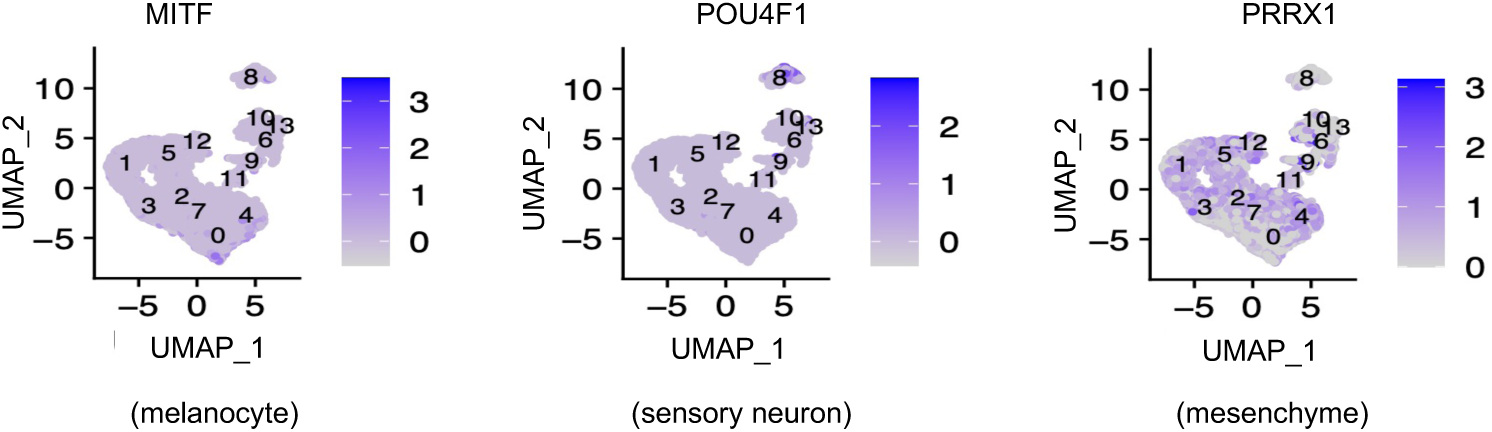
Expression of three lineage marker genes. *MITF* (melanocytes), *POU4F1* (sensory neurons), and *PRRX1* (mesenchyme), in the merged UMAP of SOX10-NL NCSCs and original NCCs.

**Supplementary Fig. 6.**
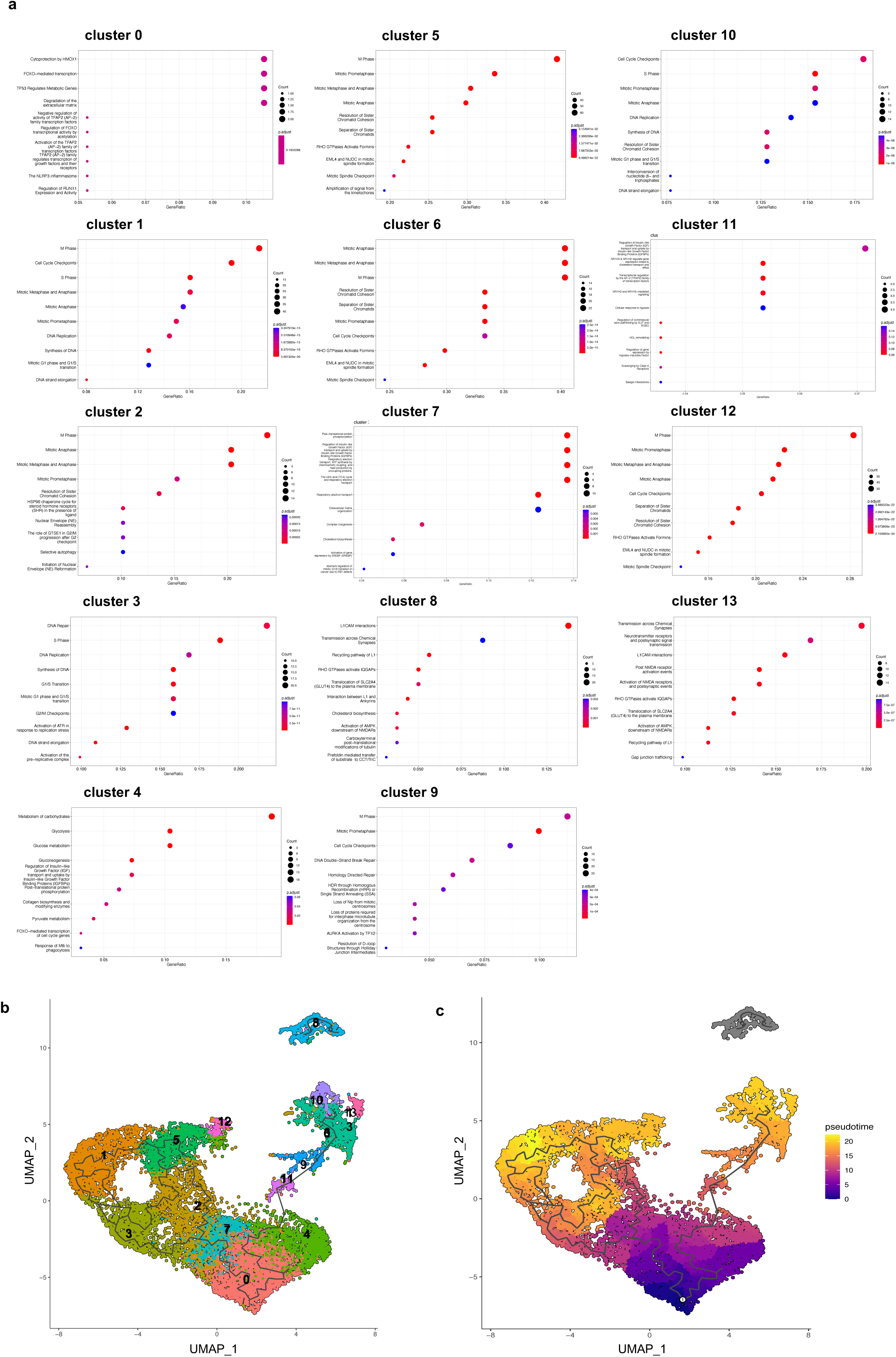
Results of scRNA-seq analysis of SOX10-tdT NCSCs. **a.** Results of reactome pathway analysis per cluster. The top ten ranked GO terms in clusters 0–13 according to gene counts are shown. The vertical items are the names of the GO terms, and the length of the horizontal graph represents the gene ratio (the percentage of total DEGs enriched in a GO term). The color depth represents the adjusted p-value. The area of the circle in the graph represents gene counts. Reactome pathway analysis was performed using the R package ‘ReactomePA’ (version 1.40.0). **b.** Result of pseudo-time analysis of the SOX10-tdT NCSCs with cluster information. The main cluster consisting of clusters 0–5 and 7 represents a loop structure, where any of the clusters 0–5 and 7 can be set as the starting point. **c.** Result of pseudo-time analysis with pseudo-time indication when cluster 0 is used as the starting point.

**Supplementary Fig. 7.**
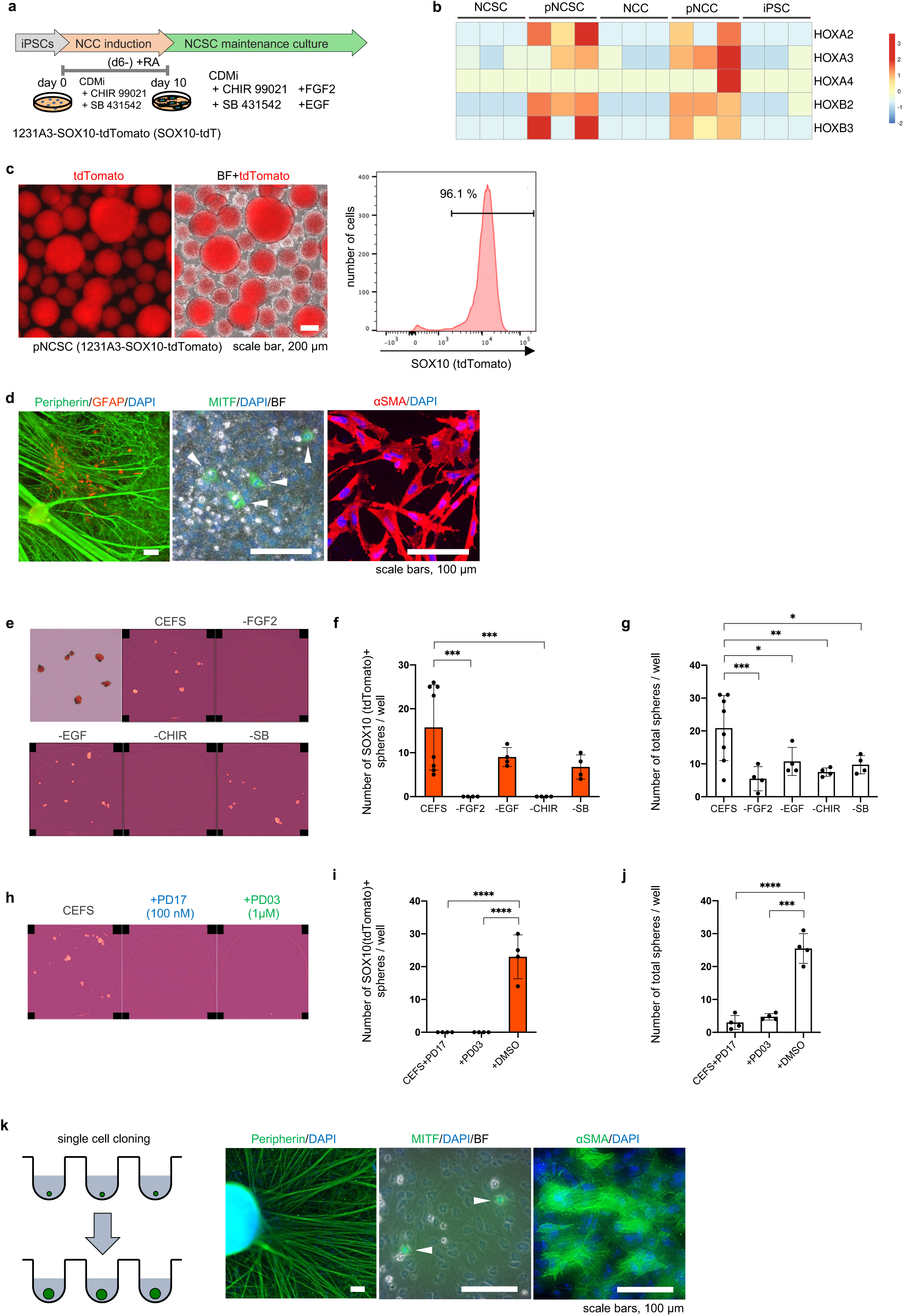
Establishment of posteriorized NCC-derived NCSCs (pNCSCs) and confirmation of the multipotency of pNCSCs at the single cell level. **a.** Schematic procedure of the induction of posteriorized NCCs and maintenance as pNCSCs using SOX10-tdT iPSCs. **b.** Heatmap presentation of the expression levels of *HOX* genes in iPSCs, NCCs, NCSCs, pNCCs, and pNCSCs (n = 3 in each cell types, biological replicates). **c.** Morphology and tdTomato fluorescence of SOX10-tdT pNCSC spheres cultured for 126 days. The SOX10+ (tdTomato+) population became dominant (approximately 90%) in established pNCSCs confirmed by FACS analysis. **d.** Differentiation of peripheral neurons (left), melanocytes (middle), and SMCs (right) from SOX10-tdT pNCSCs. **e–j.** Factor subtraction and signaling pathway inhibition experiments using SOX10-tdT pNCSCs. **e.** Representative images of each well (n = 3). Red: tdTomato fluorescence. **f, g.** Graph showing the result of factor subtraction experiments for self-renewal of pNCSCs. Means (bars), s.d. (thin lines on bars of means), and raw data (dots) are shown. **f.** The number of SOX10+ (tdTomato+) spheres formed in the wells of a 24-well plate is shown. **g.** The total sphere number in each well is shown. **h.** Representative images of the factor subtraction experiments (n = 3). **i, j.** Graph showing the result of inhibitor experiments for self-renewal of SOX10-tdT pNCSCs. Means (bars), s.d. (thin lines on bars of means), and raw data (dots) are shown. **i.** The number of SOX10+ (TdTomato+) spheres formed in the wells of a 24-well plate is shown. **j.** The total sphere number in each well is shown. **k.** pNCSC clones were differentiated into three representative NCC lineages. Schematic representation of the single cell cloning and expansion (left) and representative images of the differentiation ability of cloned pNCSCs maintained for 174 days (right). Peripheral neuron, melanocyte, and SMC differentiation were confirmed. Representative examples of the three experiments are shown. The result is summarized in Figure 3**g**. P-values were calculated using a one-way ANOVA comparison of the mean values of each condition to that of the control, followed by multiple comparisons using Dunnett’s method. *P < 0.05, **P < 0.01, ***P < 0.001, and ****P < 0.0001 indicate statistical significance.

**Supplementary Fig. 8.**
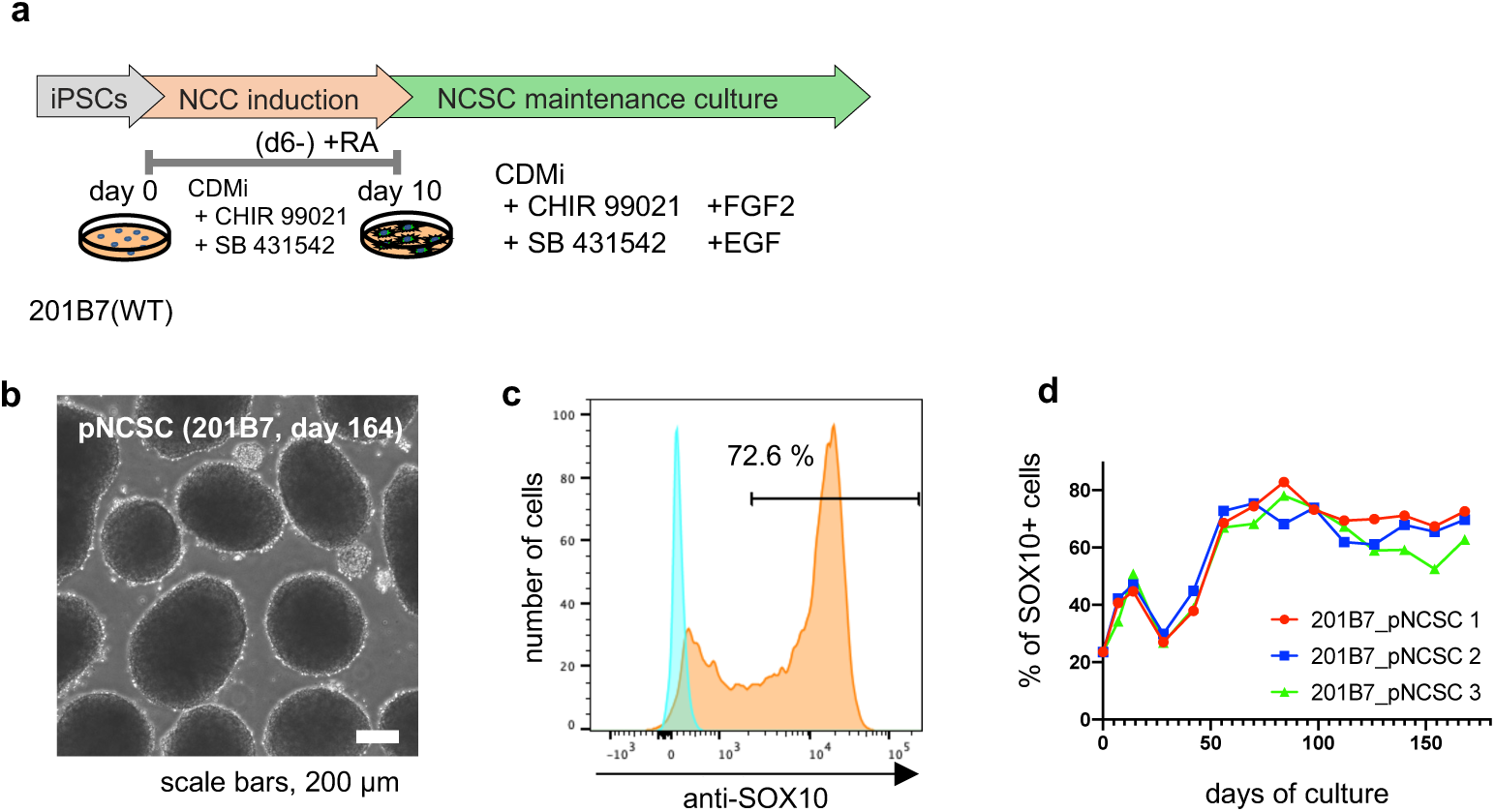
Reproducibility using 201B7 wild-type iPSCs. **a.** Schematic procedure of induction of pNCC differentiation and maintenance as pNCSCs. **b.** Bright field images of 201B7-pNCSCs cultured for 164 days. Representative images of pNCSC spheres. **c.** FACS analysis to measure the population of SOX10+ cells on day 164 from the start of maintenance culture. 201B7-pNCSCs were stained with anti-SOX10 antibody and the SOX10+ proportion was measured using FACS. **d.** Time-course changes of the SOX10+ cell proportion in the cultures from three independent experiments.

**Supplementary Table 1.** List of the antibodies used in this study.

| name of antibody | manufacture, catalog number | host | antigen | dilution used |
| --- | --- | --- | --- | --- |
| primary antibody |  |  |  |  |
| Sox10 (D5V9L) Rabbit mAb | Cell Signaling Technology, 89356S | rabbit | Sox10 | 1/150 for FACS |
| Alexa Fluor® 647 Mouse anti-Human CD271 | BD Pharmingen, cat# 560326 | mouse | NGFR (p75) | 1/5 for FACS |
| Alexa Fluor® 647 Mouse IgG1 κ Isotype Control | BD Pharmingen, cat# 557714 | mouse | - | 1/5 for FACS (Isotype control for CD271) |
| Anti-MITF antibody produced in rabbit polyclonal | SIGMA, SAB4501879 | rabbit | MITF | 1/200 for ICC<br>1/100 for IHC |
| Anti-alpha smooth muscle Actin antibody (α-SMA) | abcam, ab5694 | rabbit | α-SMA | 1/200 for ICC<br>1/500 for IHC |
| Anti-Actin, α-Smooth Muscle antibody | SIGMA, A5228 | mouse | α-SMA | 1/200 for ICC |
| Anti-Peripherin Antibody (A-3) | SantaCruz, sc-377093 | mouse | Peripherin | 1/200 for ICC |
| Anti-Peripherin antibody | abcam, ab4666 | rabbit | Peripherin | 1/500 for IHC |
| Anti-GFAP antibody | abcam, ab53554 | goat | GFAP | 1/200 for ICC |
| Anti-GFAP (GA5) antibody | Cell Signaling, #3670S | mouse | GFAP | 1/300 for ICC |
| Anti-GFAP antibody | abcam, ab7260 | rabbit | GFAP | 1/500 for ICC |
| Brn-3a (14A6) antibody | SantaCruz, sc-8429 | mouse | BRN3A | 1/50 for ICC |
| Anti-BRN3A antibody [EPR23257-285] | abcam, ab245230 | rabbit | BRN3A | 1/50 for ICC |
| Anti-Islet 1 antibody | abcam, ab20670 | rabbit | ISL1 | 1/200 for ICC |
| tdTomato Goat Polyclonal Antibody | ORIGENE, AB8181-200 | goat | tdTomato | 1/1000 for IHC |
| APC Mouse anti-CD45 | BD Pharmingen, cat# 560973 | mouse | CD45 | 1/5 for FACS |
| APC Mouse anti-CD73 | BD Pharmingen, cat# 560847 | mouse | CD73 | 1/20 for FACS |
| APC Mouse anti-CD90 | BD Pharmingen, cat# 559869 | mouse | CD90 | 1/20 for FACS |
| APC Mouse anti-CD105 | eBioscience, cat# 17-1057-42 | mouse | CD105 | 1/20 for FACS |
| APC Mouse IgG1, κ Isotype Control | BD Pharmingen, cat# 555751 | mouse | - | 1/5 for FACS (Isotype control for CD45, CD73, CD90, CD105) |
| APC Mouse anti-CD44 | BD Pharmingen, cat# 559942 | mouse | CD44 | 1/5 for FACS |
| APC Mouse IgG2b κ Isotype Control | BD Pharmingen, cat# 555745 | mouse | - | 1/5 for FACS (Isotype control for CD44) |
| APC Mouse Anti-Human HLA-DR | BD Pharmingen, cat# 340549 | mouse | HLA-DR | 1/20 for FACS |
| APC Mouse IgG2a, κ Isotype Control | BD Pharmingen, cat# 555576 | mouse | - | 1/5 for FACS (Isotype control for HLA-DR) |
| secondary antibody |  |  |  |  |
| Donkey anti-Rabbit IgG (H+L) Highly Cross-Adsorbed Secondary Antibody, Alexa Fluor™ 647 | Thermo Fisher scientific, cat # A-31573 | Donkey |  | 1/1000 for ICC |
| Donkey anti-Rabbit IgG (H+L) Highly Cross-Adsorbed Secondary Antibody, Alexa Fluor™ 546 | Thermo Fisher scientific, cat # A-10040 | Donkey |  | 1/1000 for ICC |
| Donkey anti-Rabbit IgG (H+L) Highly Cross-Adsorbed Secondary Antibody, Alexa Fluor™ 488 | Thermo Fisher scientific, cat # A-21206 | Donkey |  | 1/1000 for ICC |
| Donkey anti-Mouse IgG (H+L) Highly Cross-Adsorbed Secondary Antibody, Alexa Fluor™ 647 | Thermo Fisher scientific, cat # A-31571 | Donkey |  | 1/1000 for ICC |
| Donkey anti-Mouse IgG (H+L) Highly Cross-Adsorbed Secondary Antibody, Alexa Fluor™ 546 | Thermo Fisher scientific, cat # A-10036 | Donkey |  | 1/1000 for ICC |
| Donkey anti-Mouse IgG (H+L) Highly Cross-Adsorbed Secondary Antibody, Alexa Fluor™ 488 | Thermo Fisher scientific, cat # A-21202 | Donkey |  | 1/1000 for ICC |
| Donkey anti-Goat IgG (H+L) Cross-Adsorbed Secondary Antibody, Alexa Fluor™ 488 | Thermo Fisher scientific, cat # A-11055 | Donkey |  | 1/1000 for ICC |
| ChIP-seq analysis |  |  |  |  |
| Anti-SOX10 antibody | abcam, ab272074 | Rabbit | SOX10 |  |
| Rabbit IgG, monoclonal, Isotype Control | abcam, ab210849 | Rabbit | - | control for ChIP-seq |

